# Lasp family proteins generated by teleost-specific genome duplication

**DOI:** 10.64898/2026.10.05.756842

**Authors:** Yuhang Xie, Kouta Morimoto, Ryota Tsunakawa, Yasushige Mori, Hina Awaya, Sakura Haruta, Yuki Fujita, Tamami Chokki, Ayako Nakayama, Yuu Usui, Yu Katsuyama, Asako G Terasaki

## Abstract

Two rounds of whole-genome duplication (2R-WGD) in the vertebrate lineage are thought to have driven gene diversification. Mammals have four genes encoding five nebulin superfamily proteins (*LASP1*, *LASP2/NEBL*, *NEB*, and *NRAP*) hypothesised to have arisen from invertebrate *LASP* via WGD. Here, we examined the evolutionary fate of the nebulin superfamily after an additional WGD, teleost-specific genome duplication (TGD). Genome and transcriptome analysis of zebrafish (*Danio rerio*) and medaka (*Oryzias latipes*), of which the complete genomes were recently determined, showed only lasp family genes retained duplicate copies. Zebrafish retained two *LASP2/NEBL*-derived ohnologs; one (*LASP2L*) had been misidentified as a nebulette gene in the Zebrafish Information Network (ZFIN) despite lacking nebulette-specific exons. Another (*LASP2/NEBL*) is newly identified in this study because it lies in a newly determined region of the latest genome assembly. Contrastingly, medaka retained two *LASP1* ohnologs and one *LASP2/NEBL*. The duplicated lasp family proteins showed overlapping and specific tissue expression patterns. We confirmed all three zebrafish lasp proteins bound actin and co-localized with actin-rich structures when ectopically expressed in cultured fibroblasts. This supports that TGD-derived lasp duplicates underwent lineage-specific retention and expression divergence in teleosts whilst conserving core actin-binding properties.

## 1 Introduction

Two rounds of whole-genome duplication (2R-WGD) have occurred in the vertebrate lineage, and logically, generating four paralogous (reviewed by Altschmied et al.). Most duplicated genes generated by WGD (ohnologs) underwent “non-functionalization” (pseudogenization) through the loss of promoter or coding-region function, due to the accumulation of mutations. Others were retained through mechanisms, including “neofunctionalization” and “subfunctionalization”, where they specialize through the partitioning of ancestral gene functions (reviewed by Kuzmin et al. 2022).

In Mammalia and Aves, four genes encode five nebulin-repeat proteins of the nebulin superfamily: lasp-1 (LIM and SH3 protein 1) transcribed from *LASP1* (gene symbol used in human); lasp-2 (LIM and SH3 protein 2, also called LIM-nebulette) and nebulette from *NEBL;* nebulin from *NEB; and* N-RAP (nebulin-related-anchoring protein) from *NRAP* (Pappas et al. 2011). We previously identified unique alternative splicings of *NEBL* in Mammalia, Aves, Amphibia, and Actinopterygii (Terasaki et al., 2006; Fujita et al., 2021) and conserved among vertebrate species; we refer to this as “*LASP2/NEBL”.* All these members from the nebulin superfamily have actin-binding activity and localize to various actin-containing structures (see Fig. 1 of Fujita et al. 2021). In mammals and avians, lasp-1 is expressed in various non-muscle tissues and localizes in actin-rich subcellular regions such as lamellipodia, filopodia, and focal adhesions (Valé Rie Schreiber et al., 1998; Chew, 2002). Lasp-2 is highly expressed in the brain, whereas nebulette is specifically expressed in cardiac muscles (Moncman & Wang, 1995; Terasaki et al., 2004). Cellular localization of lasp-2 is similar to lasp-1, except in cardiac muscles, where lasp-1 shows no apparent specific localization and lasp-2 is restricted to Z-lines (Zieseniss et al. 2008). NRAP localizes in intercalated discs, which connect cardiomyocytes during repeated contraction, and myotendinous junctions. These overlapping and distinct localization patterns suggest that nebulin superfamily proteins cooperate in regulating the actin cytoskeleton.

**Fig. 1.**
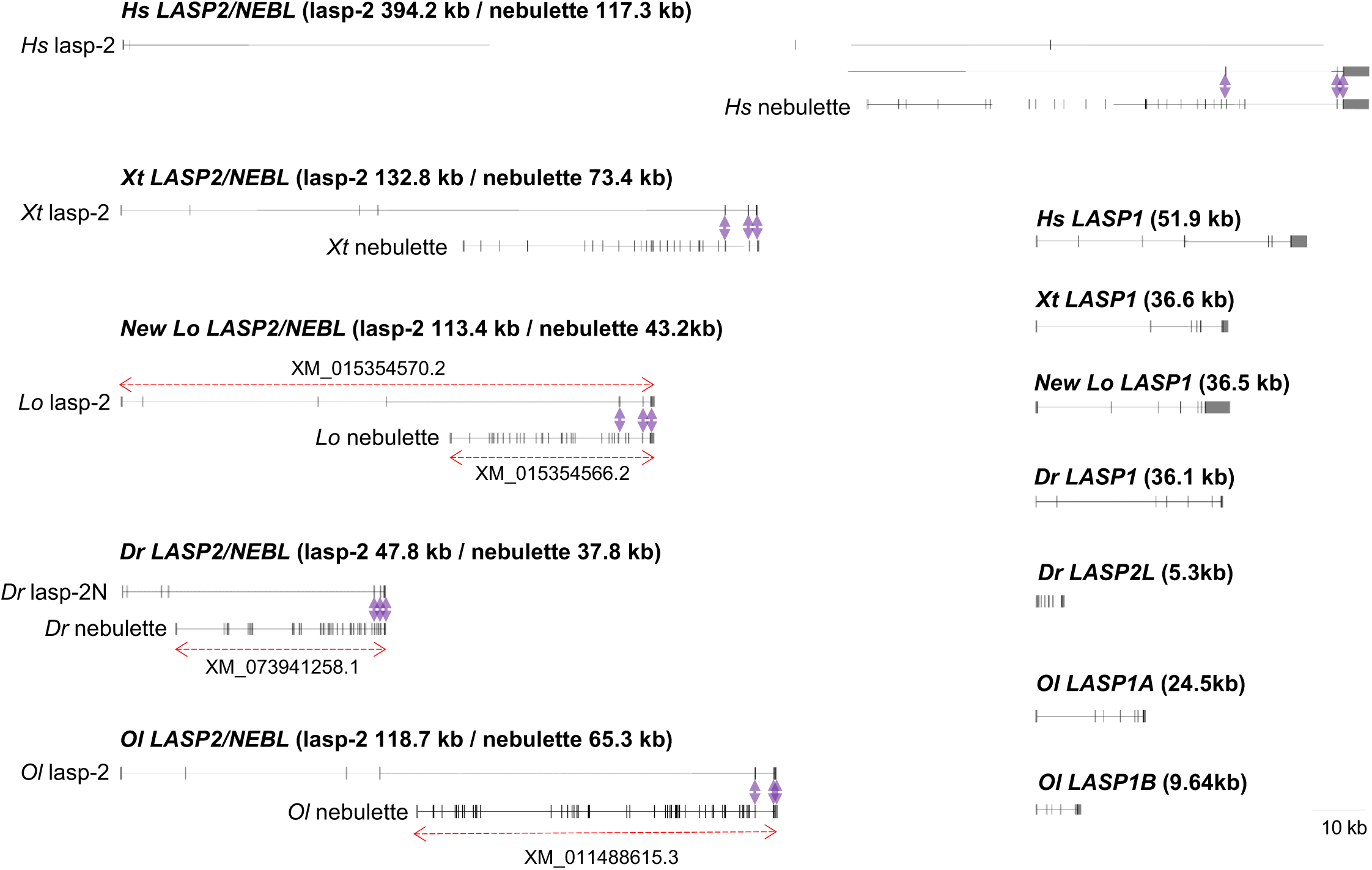
Gene structures of lasp family proteins of teleost. Intron lengths and overall gene lengths of *LASP1*, *LASP2/NEBL*, and *LASP2L* of human (*Hs*), tropical frog (*Xt*), spotted gar (*Lo*), zebrafish (*Dr*), and medaka (*Ol*). The horizontal lines depict genomic regions of the genes; exons are indicated vertically. In *LASP2/NEBL*, genomic regions coding lasp-2 and nebulette are indicated, respectively. Double arrows indicate shared exons between lasp-2 and nebulette. The large genomic region of *Hs LASP2/NEBL* is illustrated across two rows. *LASP1* and *LASP2/NEBL* of human and tropical frog were cited from our previous study (Fujita et al., 2022), and *LASP1* and *LASP2/NEBL* of spotted gar were updated using a new genome assembly.

Based on domain structure, our previous study proposed that the four vertebrate nebulin superfamily genes arose from a single ancestral gene, as all members contain nebulin repeats together with the LIM domain and/or the SH3 domain (Terasaki et al., 2006). We subsequently identified lasp (*LASP*), which has similar domain structure to vertebrate *LASP1* and *LASP2/NEBL*, in the invertebrate sea squirt (*Ciona intestinalis*) (Terasaki et al. 2008) and revealed that invertebrates within Filozoa, except for Placozoa, have at least one *LASP* (Fujita et al. 2021a). Additionally, synteny and exon-intron boundaries indicated that vertebrate *LASP1* and *LASP2/NEBL* evolved from ancestral *LASP* (Fujita et al. 2021). The actin-binding activity of lasp of sea squirt, fruit fly (*Drosophila melanogaster*), and nematode (*Caenorhabditis elegans*) was confirmed (Terasaki et al. 2008; Suyama et al. 2009; Fujita et al. 2021). Thus, the four vertebrate nebulin superfamily genes may have undergone “neofunctionalization” and “subfunctionalization” accompanying the duplication of nebulin repeats loss of the LIM or SH3 domain after 2R-WGD.

Teleostei (modern bony fishes) is a group within Actinopterygii (ray-finned fish) and underwent an additional round of whole-genome duplication, the teleost-specific genome duplication (TGD), after divergence from tetrapods (reviewed by Glasauer and Neuhauss 2014). TGD is considered to have contributed to the remarkable species diversity of teleosts (Inoue et al. 2015; reviewed by Pasquier et al. 2017). Although more than 200 actin-binding proteins have been identified in mammals and avians, their TGD-derived duplicates and functions in teleosts remain poorly studied. Moreover, it is unclear whether duplicated actin-binding proteins underwent “neofunctionalization” or “subfunctionalization” in teleosts.

We previously identified lasp-1, lasp-2, and nebulette genes and transcripts in spotted gar (*Lepisosteus oculatus, Lo*), a member of Holostei that diverged before TGD and therefore retains more ancestral genomic features (Fujita et al. 2021a). In the current study, we analyzed gene duplication of nebulin superfamily genes in zebrafish (*Danio rerio, Dr*), which has a complete genome assembly and the integrated database Zebrafish Information Network (ZFIN; https://zfin.org/) (Kimmel et al. 1995). We also analyzed medaka (*Oryzias latipes, Ol*), another teleost of which a complete genome assembly has determined. Using PhyloFish database (http://phylofish.sigenae.org/index.html) (Pasquier et al. 2016), an mRNA database of TSA (Transcriptome Shotgun Assembly) generated by de novo assembly from SRA (Sequence Read Archive), together with genome-predicted RefSeq sequences, we confirmed that only lasp family genes were duplicated in these two model organisms. Gene structure, domain structures, synteny, tissue expression patterns of lasp genes of zebrafish, medaka, and spotted gar suggested duplication of *LASP2/NEBL* occurred in zebrafish whereas duplication of *LASP1* occurred in medaka. To investigate the functional diversity of zebrafish lasp family proteins, we analyzed actin-binding activity and localization of ectopically expressed proteins in cultured cells.

For cross-species comparisons, gene names are italicized and capitalized following human nomenclature (e.g., *Dr LASP1*), whereas mRNA and protein names are lowercase (e.g., *Dr* lasp-1).

## 2 Results

### 2.1 Only lasp family protein genes were duplicated among teleost nebulin superfamily

We recently identified genes and transcripts of *Lo LASP1* and *Lo LASP2/NEBL* of spotted gar (Fujita et al., 2022); however, the previous genome assembly of spotted gar (fLepOcu1.hap2) contained gaps, so we reanalyzed the gene structures of *Lo LASP1* and *Lo LASP2/NEBL* using the updated genome assembly (fLepOcu2.hap2). The genomic region of *Lo LASP1* changed from 36.8 kb to 36.5 kb and *Lo LASP2/NEBL* changed from (lasp-2 116.7 kb / nebulette 47.1kb) to (lasp-2 113.4 kb / nebulette 43.2kb), without changes in the number of exons of each transcript and the position of nebulette exons inserted between lasp-2 exons (Fig. 1). We updated only one amino acid of *Lo* lasp-1 following the new genome assembly (supplemental data 1).

Using amino acid sequences of *Lo* lasp-1 and *Lo* lasp-2 as queries, we searched for lasp family proteins of zebrafish and medaka in Phylofish and detected three transcripts encoding complete LIM domain, nebulin repeats, and SH3 domain (Table S1). In zebrafish, transcripts for three lasp family proteins (FDR_LASP1.1.1, FDR_NEBL.2.2, and FDR_LNEBL.2.2) were identified from Phylofish. FDR_LASP1.1.1 was identical to NM_212636.1 on chromosome 3 in the zebrafish genome assembly (GRCz12tu) and annotated as “lasp-1” (ZFIN gene ID ZDB-GENE-030131-1936; NCBI gene ID 323216). FDR_NEBL.2.2 was identical to “nebulette”(ZFIN gene ID ZDB-GENE-050320-151; NCBI gene ID 541446) on chromosome 2. However, FDR_LNEBL.2.2 was absent from ZFIN and only partially overlapped an uncharacterized gene (ID 137489732) on chromosome 24. We further identified a longer zebrafish transcript (FDR_LOC100707579.2.2) that shares 3’ sequences with FDR_LNEBL.2.2 in the same genomic region. Three medaka lasp-family transcripts (LOL_LOC101156576.3.3, LOL_LOC101165512.1.1, and LOL_LNEBL.1.1) were also identified from Phylofish, of which corresponding RefSeq RNAs in the medaka genome assembly (Olatipes_Hd-rR_3.1) were found on chromosomes 8, 19, and 20, respectively. LOL_LNEBL.1.1 shares 3’ sequences with LOL_LOC101166998.3.8 from Phylofish, similar to FDR_LNEBL.2.2/FDR_LOC100707579.2.2. The original LOL_LOC101166998.3.8 (9008 bp) contains a sequence with multiple stop codons, likely derived from SRA of immature mRNA because this region was absent from two medaka EST clones (olbrno43_l05 and olbrno18_a24). Therefore, an intron-removed version of LOL_LOC101166998.3.8 was used for analysis (Table S1), Together, these data showed that one of three lasp family genes from each of the two species is *LASP2/NEBL* (see section 2.2).

Using N-RAP and nebulin of human (*Homo sapiens, Hs*) as queries, we identified transcripts of N-RAP (BLO_contig_017979, LIM domain and nebulin repeats) and nebulin (BLO_NEB.1.7, nebulin repeats and SH3 domain) of spotted gar, whose tissue expression patterns are similar to those in humans (Table S1); these transcripts corresponded to RefSeq RNA (XM_006630835.3) on chromosome 10 and RefSeq RNA (XM_069196485.1) on chromosome 12 in the spotted gar genome assembly (fLepOcu1.hap2). We also confirmed that the zebrafish N-RAP transcript (FDR_NRAP.1.1) was identical to “nrap” in ZFIN (ZFIN gene ID: ZDB-GENE-060616-224). All nebulin transcripts of zebrafish identified in Phylofish partially matched XM_073912668.2, a predicted *neb* transcript on chromosome 9 encoding 171 nebulin repeats and an SH3 domain (ZFIN gene ID: ZDB-GENE-041111-216). Similarly, medaka had single N-RAP and nebulin genes, along with their corresponding transcripts (Table S1). No additional transcripts containing nebulin repeats, other than those described above, were identified in the genome assemblies of spotted gar, zebrafish, and medaka. Thus, spotted gar has the same four nebulin superfamily genes as humans (*LASP1*, *LASP2/NEBL*, *NRAP*, and *NEB*); moreover, among these genes, only the lasp family was duplicated in zebrafish and medaka.

### 2.2 Gene structures of three lasp family proteins of zebrafish and medaka

We compared the exon–intron boundaries and intron lengths of the three transcripts of lasp family proteins from zebrafish and medaka with the *LASP1* and *LASP2/NEBL* transcripts from spotted gar, tropical frog, chicken, and human analyzed in our previous study Fujita et al. 2021 ; Fig. 1). *Dr LASP1* transcript (FDR_LASP1.1.1) consists of seven exons, and its gene structure and genomic length are similar to those of *LASP1* in tetrapods and spotted gar. However, FDR_NEBL.2.2, annotated as “nebl” in ZFIN, spans a much smaller region and lacks the nebulette-specific exons conserved in tetrapods and Actinopterygii (Fujita et al. 2021). Thus, FDR_NEBL.2.2 does not encode nebulette. Instead, the gene structure defined by FDR_LNEBL.2.2 and FDR_LOC100707579.2.2 resembles *Lo LASP2/NEBL* (Fig. 1; see also Fig. 3B). We therefore designated FDR_LNEBL.2.2 as Dr lasp-2N (lasp-2-new) and FDR_LOC100707579.2.2 as *Dr* nebulette, neither of which is annotated in ZFIN. FDR_NEBL.2.2, formerly annotated as nebulette, was designated *Dr* lasp-2L (lasp-2-like), because it was shown to be a WGD-derived duplicate (detailed in following sections). These designations are used hereafter.

Unlike the transcripts of lasp-2 (BLO_LNEBL.1.1) and nebulette (BLO_LOC100707579.1.1) of spotted gar in Phylofish, which closely matched their RefSeq sequences (XM_015354570.2 and XM_015354566.2), the transcripts of nebulette and lasp-2 of zebrafish and medaka in Phylofish differ from the corresponding NCBI RefSeq predictions (Fig. 1, Table S1). A BLAST search of zebrafish nebulette (FDR_LOC100707579.2.2) against GRCz12tu revealed the transcript possesses a 3,879 bp coding sequence (CDS) comprising 39 exons, and encodes 28 nebulin repeats and one SH3 domain as determined by the SMART (Table S1). Contrastingly, the corresponding RefSeq prediction (XM_073941258.1) contains a 6,192 bp CDS with 62 exons, encoding 48 nebulin repeats and one SH3 domain. Thus, 23 exons (exons 3–5, 16–28, and 33–39) of XM_073941258.1 are absent from FDR_LOC100707579.2.2. Similarly, compared with Olatipes_Hd-rR_3.1, medaka nebulette (LOL_LOC101166998.3.8) contains a 6,030 bp CDS comprising 54 exons, and encodes 50 nebulin repeats and one SH3 domain (Table S1). The corresponding RefSeq prediction (XM_011488615.3) also contains a 7,569-bp CDS with 14 additional exons. Additional exons of both zebrafish and medaka encode nebulin repeats. In both species, RefSeq predicted only nebulette transcripts, although the first four exons of *Dr* lasp-2N (FDR_LNEBL.2.2) and *Ol* lasp-2 (FOL_LNEBL.2.2) are located upstream of these RefSeq transcripts (Fig. 1). The gene structure of *LASP2/NEBL* in both teleost species was highly similar to that of *LASP2/NEBL* of tetrapods and spotted gar (Fig. 1 and 3A). Spotted gar and zebrafish *LASP2/NEBL* contain two nebulette-specific exons, corresponding to human exons 25 and 26, whereas *Ol LASP2/NEBL* lacks these exons, as in chicken and tropical frog (Fujita et al. 2021).

### 2.3 Syntenic analysis indicates that zebrafish *LASP2/NEBL* and *LASP2L* and medaka *LASP1A* and *LASP1B* were generated by WGD

Using physical mapping, we analyzed the syntenic conservation among zebrafish and medaka, which underwent TGD, and spotted gar, which did not. Using the Comparative Genome Viewer, we compared spotted gar chromosome 28 containing *Lo LASP1* and chromosome 6 containing *Lo LASP2/NEBL* with all zebrafish and medaka chromosomes. Consistent with the retention of duplicated TGD-derived chromosomes in teleosts (Braasch et al., 2016), spotted gar chromosome 28 exhibited the highest synteny with zebrafish chromosome 3, harboring *Dr LASP1,* and chromosome 6 (Fig. 2A). Similarly, spotted gar chromosome 28 exhibited synteny with medaka chromosomes 8 harboring *Ol LASP1A* and 19 harboring *Ol LASP1B*. Spotted gar chromosome 6 exhibited the highest synteny with part of zebrafish chromosomes 2 harboring *Dr LASP2L* and 24 harboring *Dr LASP2/NEBL* (Fig. 2A).

**Fig. 2.**
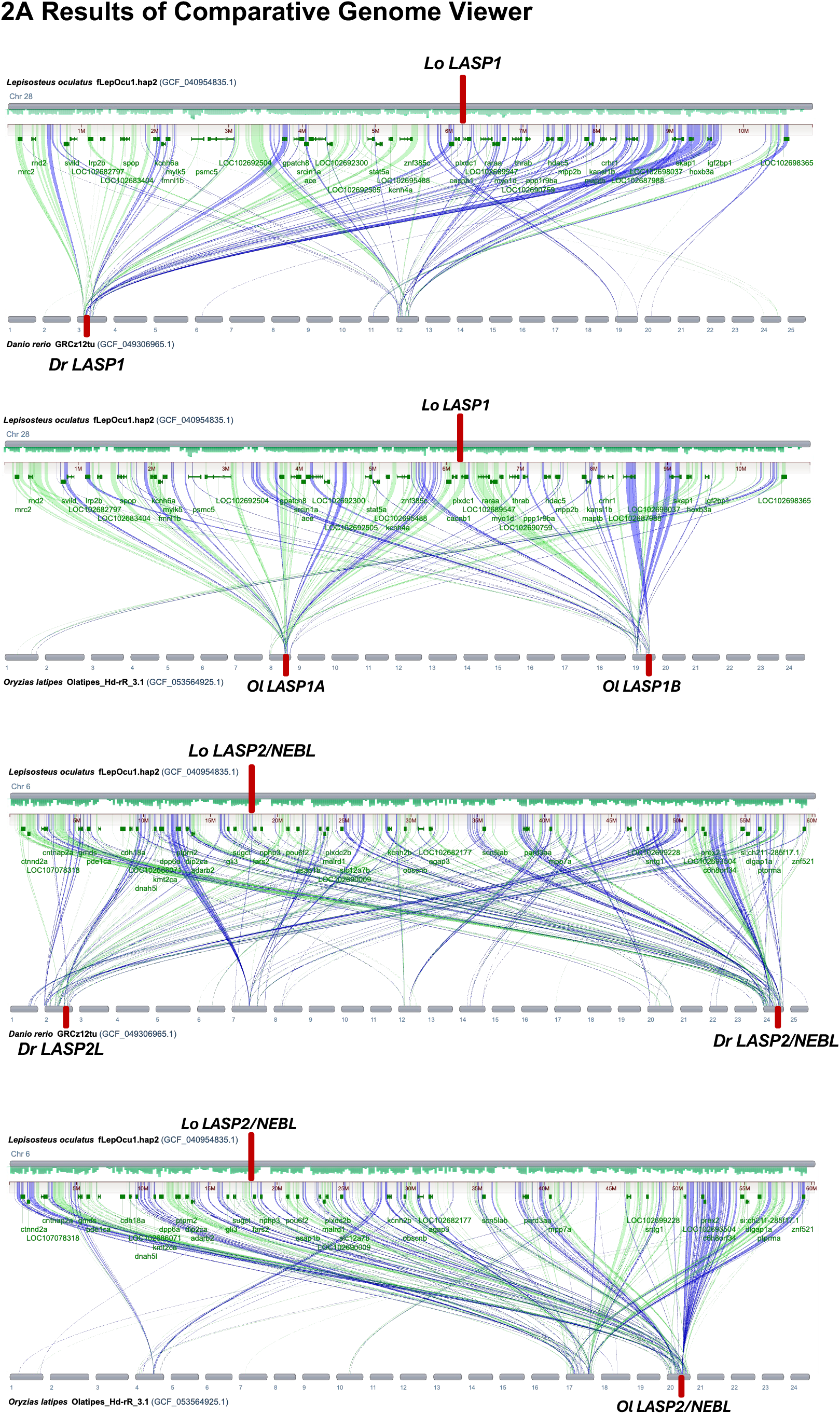
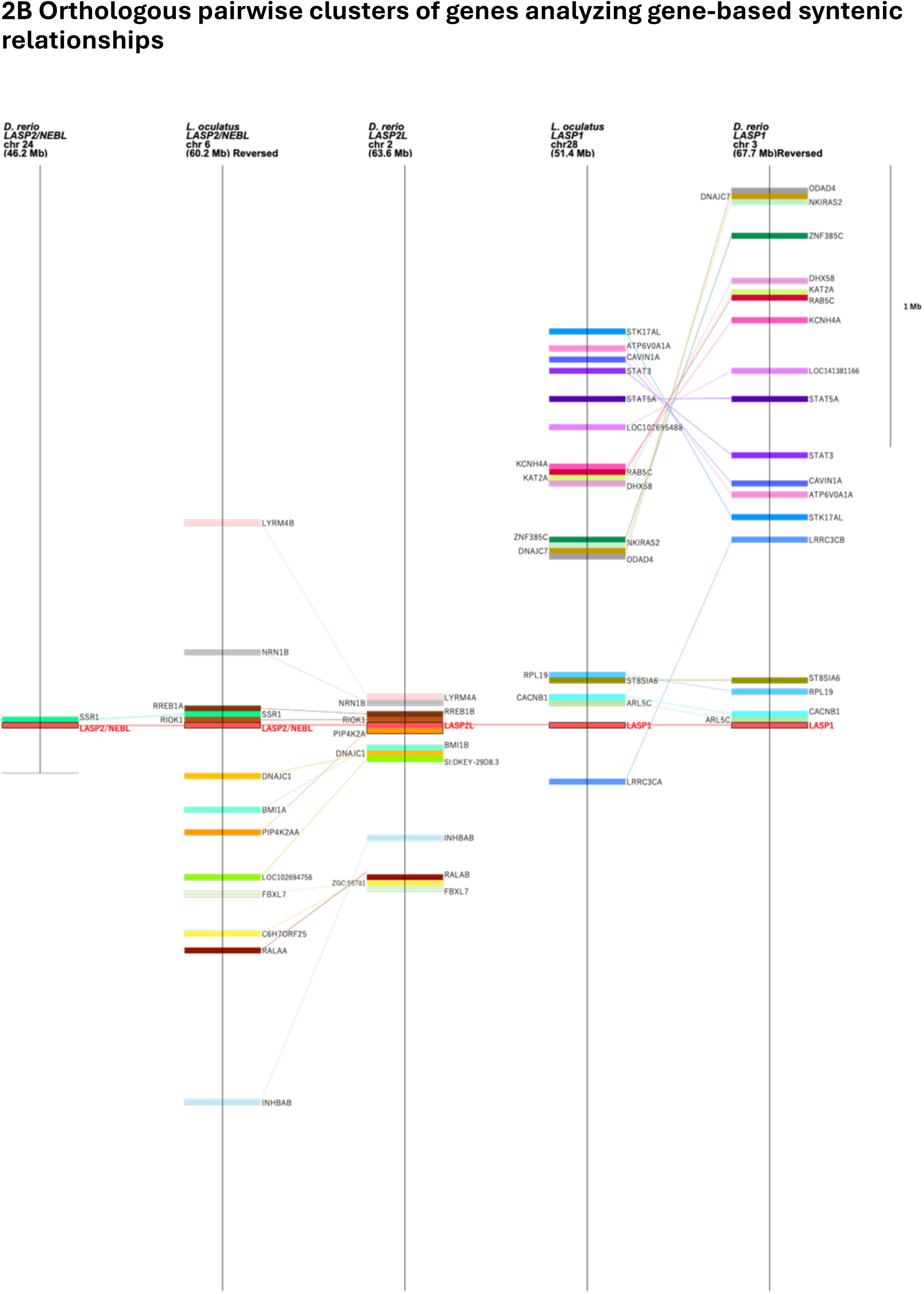
Synteny analysis lasp family proteins of spotted gar and two teleost species. A. Results of Comparative Genome Viewer analyzing DNA-based syntenic relationships between spotted gar chromosomes 6 (containing *LASP2/NEBL*) or 28 (containing *LASP1*) and all chromosomes of zebrafish and medaka, with a minimum alignment length of 10,000 bp. Locations of genes of lasp family proteins are indicated by red vertical lines. B. Orthologous pairwise clusters of genes analyzing gene-based syntenic relationships between spotted gar and zebrafish. Genes on vertical lines are located within a 2-Mb distance from genes of lasp family proteins. Supplemental Table 2 presents the genes used in the syntenic analysis.

We further compared genes surrounding lasp family loci in spotted gar and zebrafish using BLASTP (Fig. 2B). Most spotted gar genes were predicted as XM records (predicted mRNA), although many zebrafish genes were represented by NM records (experimentally identified mRNA). We therefore confirmed the expression of all genes used in the synteny analysis (Table S2). Many homologous genes surrounding *Lo LASP2/NEBL* on chromosome 6 were mapped near *Dr LASP2L* on chromosome 2, and their gene order was conserved (Fig. 2B). However, no genes homologous to those near *Lo LASP1* were found surrounding *Dr LASP2L*. Conversely, many genes neighbouring *Lo LASP1* had homologs near *Dr LASP1*.

### 2.4 Teleost lasp family proteins have conserved domain structure but duplicated lasp-2 of zebrafish lost the third nebulin repeat

All vertebrate LASP genes analyzed in this study shared a similar structure comprising seven exons (Fig. 1). We also compared the exon-intron boundaries and domain structure of three newly identified zebrafish and medaka lasp family proteins with those previously analyzed in other species, including spotted gar (Fujita et al. 2021) (Fig. 3A). No differences were observed in the first four exons—encoding the LIM domain, the first nebulin repeat, and most of the second nebulin repeat—or the seventh exon encoding the entire SH3 domain. Although lasp-2 mRNAs of other organisms possess a third nebulin repeat encoded by the fifth exon, zebrafish lasp-2N and lasp-2L lack this repeat (described later). Additionally, the nebulette exon insertion site splitting the second nebulin repeat (red arrowheads in Fig. 3A) was conserved among *Hs* lasp-2, *Gg* lasp-2, *Xt* lasp-2, *Lo* lasp-2, *Dr* lasp-2N, and *Ol* lasp-2. *Hs* lasp-2, *Lo* lasp-2, and *Dr* lasp-2N also shared a second insertion site containing two nebulette-specific exons (yellow arrowheads in Fig. 3A; Fig. 3B). *Ol* lasp-2 lacked these exons, as did *Xt* lasp-2 and *Gg* lasp-2 in our previous study (Fujita et al. 2021) (Fig. 3B).

**Fig. 3.**
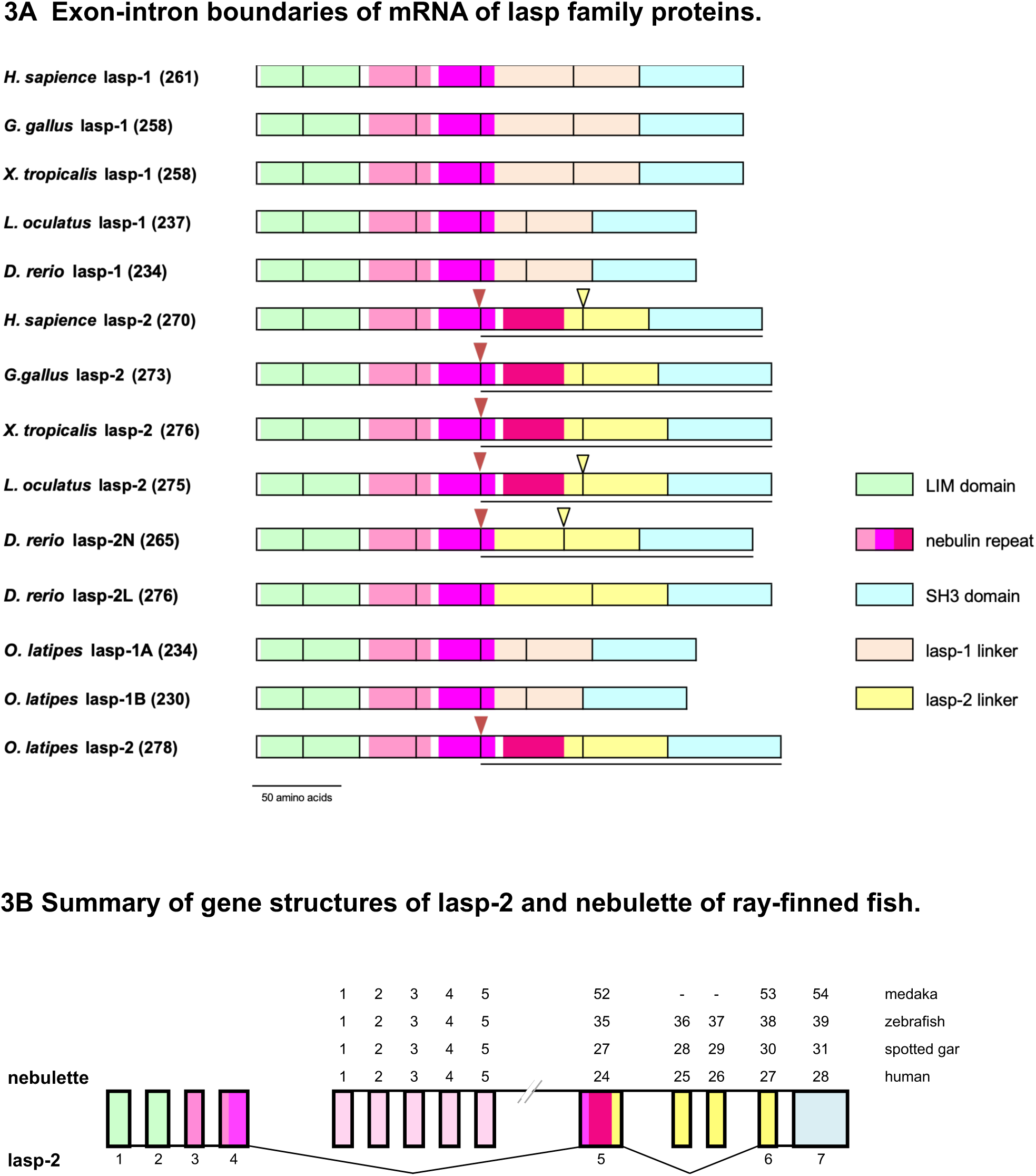
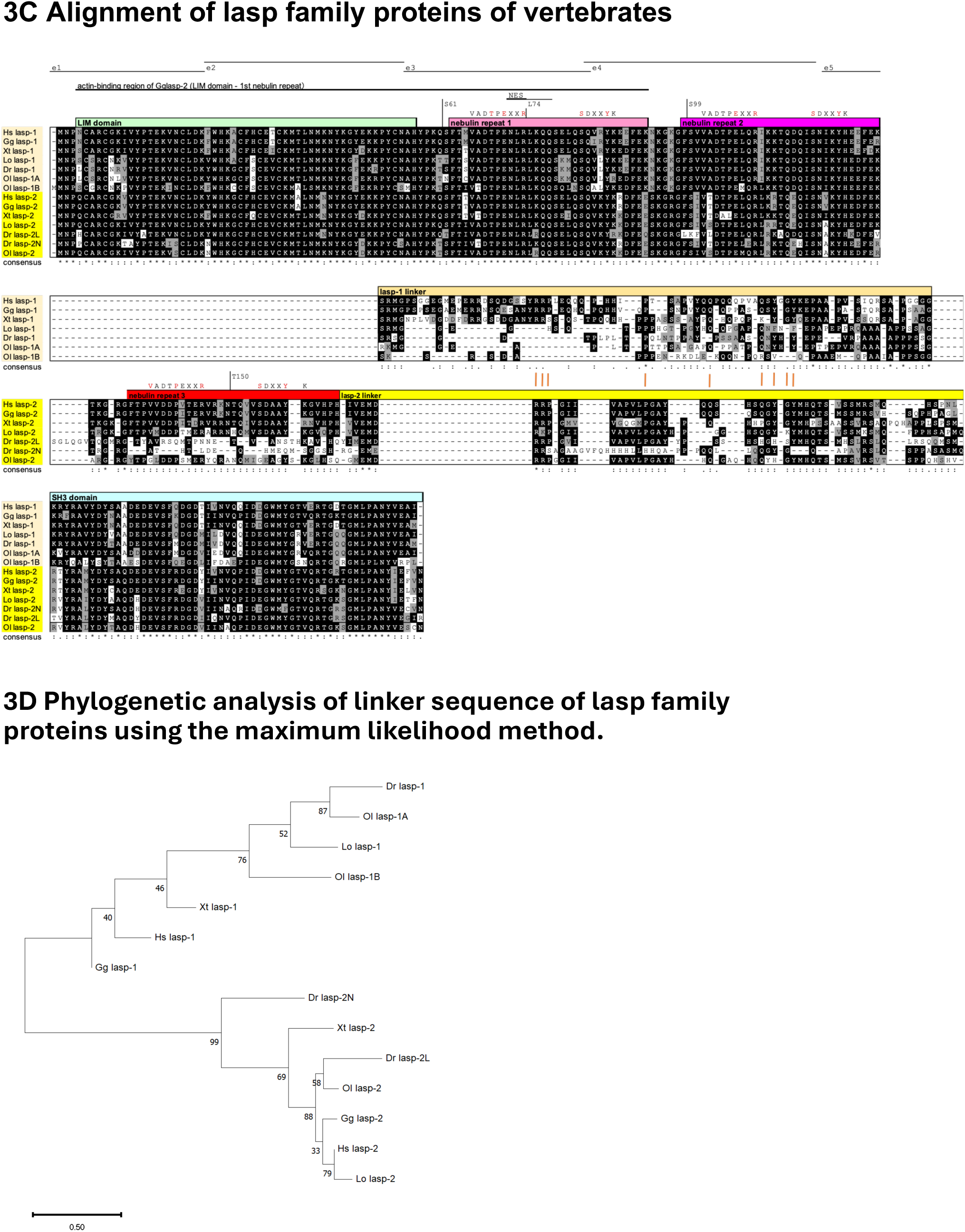
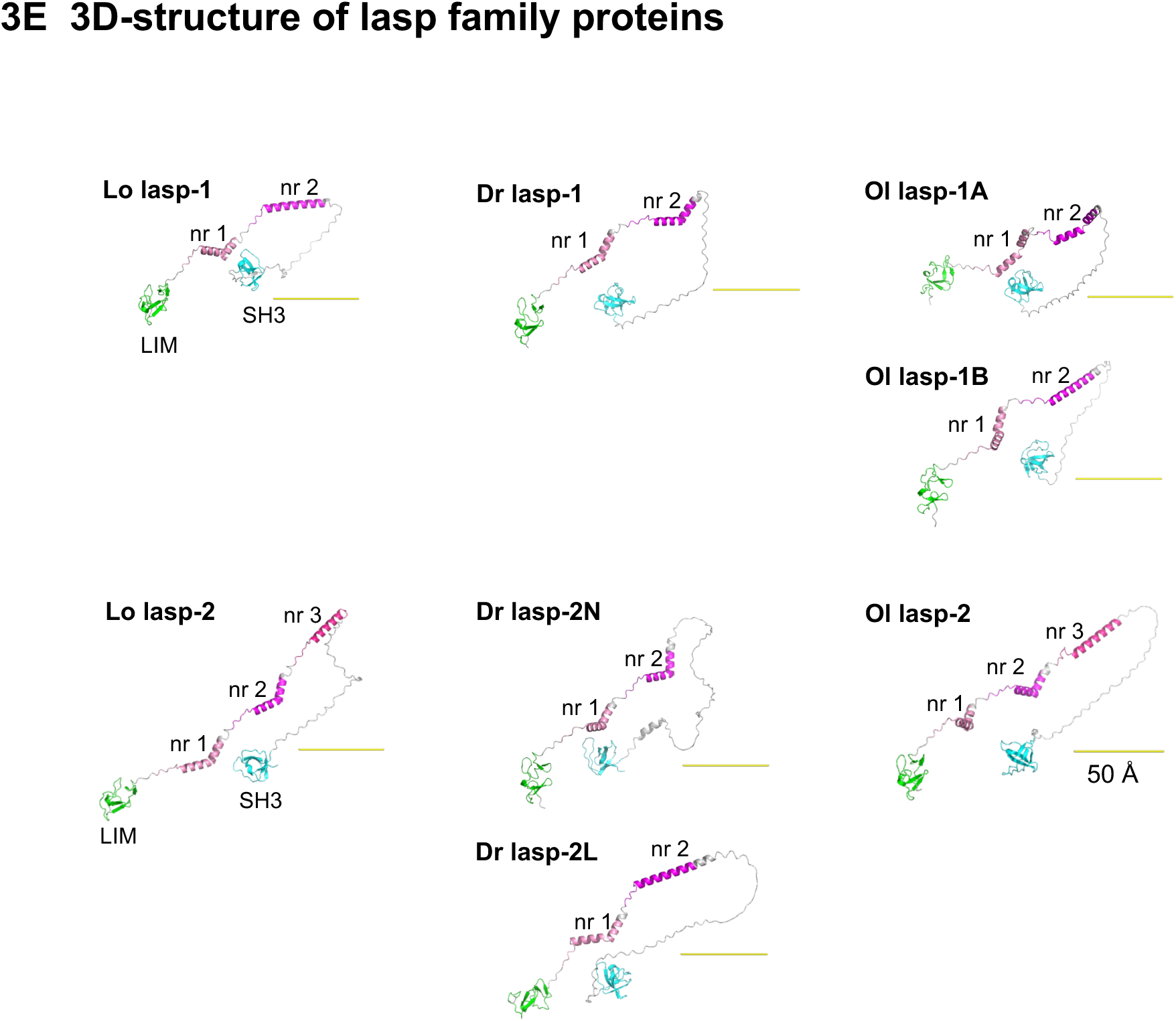
Comparison of domain structure, sequence, and 3D structure of lasp family proteins of ray-finned fish. A. Exon-intron boundaries of mRNA of lasp family proteins. Colored regions correspond to the LIM domain (green), nebulin repeats (pink, magenta, and red), SH3 domain (blue), and conserved linker sequences of vertebrate lasp-1 and lasp-2 (pale orange and yellow, respectively). Red arrowheads in *Hs* lasp-2, *Gg* lasp-2, *Xt* lasp-2, *Lo* lasp-2, *Dr* lasp-2 and *Ol* lasp-2 indicate sites where nebulette exons were inserted, and yellow arrowheads indicate additional insertion sites of *Hs* lasp-2, *Lo* lasp-2, and *Dr* lasp-2. Underlined exons are shared with nebulette. B: Summary of gene structures of lasp-2 and nebulette of ray-finned fish. As indicated in our previous study (Fujita et al. 2022), human lasp-2 mRNA consists of 7 exons and shares exons 5–7 with nebulette mRNA, which has 28 exons. In the present study, lasp-2 and nebulette of spotted gar, zebrafish, and medaka were revealed to have structures similar to those of humans. However, the total number of exons of nebulette and the insertion of two additional nebulette-specific exons between exon 5 and exon 6 of lasp-2. C. Multiple sequence alignment of the LIM domain (green), the first nebulin repeat (pink), the second nebulin repeat (magenta), and the third nebulin repeat (red) of zebrafish and medaka lasp family proteins with previously analyzed vertebrate lasp family proteins. Lines above the alignment indicate positions of exons conserved among lasp family proteins analyzed in this study; the region essential for actin-binding activity of Gg lasp-2 analyzed in our previous study (Nakagawa et al., 2009; Fujita et a;, 2022) is also indicated with a line. S61 and S99 indicate phosphorylation sites, and L74 is an essential amino acid in the NES reported in mammalian lasp-1. SDXXYK is a consensus sequence of the nebulin repeat, and VADTPEXXR is a newly identified consensus sequence revealed in our previous study (Fujita et al. 2021). Conserved residues are indicated in black and similar residues in light gray. Asterisks (*) under sequences indicate amino acids completely conserved in all species. Colons (:) indicate amino acids showing 50%–99% identity or 50%–100% similarity, and periods (.) indicate 30%–49% similarity between species. Orange vertical lines indicate homologous amino acids of linker sequences between lasp-1 and lasp-2. D. Phylogenetic analysis of linker sequence of lasp family proteins using the maximum likelihood method. Bootstrap values are shown for each node. The tree is drawn to scale, with branch lengths indicative of the number of substitutions per site. E. 3D-structure of lasp family proteins of spotted gar, zebrafish and medaka (*Lo* lasp-1, *Lo* lasp-2, *Dr* lasp-1, *Dr* lasp-2N, *Dr* lasp-2L, *Ol* lasp-1A, *Ol* lasp-1B, *Ol* lasp-2) predicted using the AlphaFold server. Coloring of the LIM domain (green), the first nebulin repeat (pink), the second nebulin repeat (magenta), the third nebulin repeat (red), and the SH3 domain (green) is used for better visualization. An α-helix represents each nebulin repeat, and the bent regions found in some nebulin repeats.

Sequence alignment of lasp family proteins revealed strong conservation in the LIM domain and first nebulin repeat, which make up the actin-binding region identified in our previous study (Nakagawa et al. 2009) (Fig. 3C), as well as the second nebulin repeat. The SDXXYK motif, originally identified in nebulin (Labeit and Kolmerer 1995), was observed as SQXXYK in the first nebulin repeat and as SNXXYH in the second. Another consensus sequence, VADTPEXXR, previously identified (Fujita et al. 2021), was also highly conserved. The SH3 domain was highly conserved across all lasp family proteins (Fig. 3C). In contrast, the region corresponding to the third nebulin repeat of *Dr* lasp-2N and *Dr* lasp-2L showed lower homology to *Lo* lasp-2 and tetrapods, indicating loss of the canonical repeat sequence. The same region in *Ol* lasp-2 also has low similarity to the canonical third nebulin repeat, but was identified as a third nebulin repeat by SMART (Fig. 3A). Alignment of lasp-1 linkers showed more diverse sequence variation in ray-finned fishes than in tetrapods. In particular, *Ol* lasp-1B showed lower similarity to *Lo* lasp-1, *Dr* lasp-1, and *Ol* lasp-1A. Ray-finned fish lasp-2 linkers were generally conserved relative to tetrapod lasp-2, except for *Dr* lasp-2N. Although lasp-1 and lasp-2 linker sequences are highly divergent, several locally conserved residues remained (orange vertical lines), suggesting a shared evolutionary origin (Fig. 3C).

As shown in Fig. 3D, the phylogenetic tree of the third nebulin repeat and linker separated lasp-1 and lasp-2 into distinct clades. Within the lasp-1 clade, the clustering pattern is generally aligned with the established species phylogeny. Tetrapod and ray-finned fish sequences generally formed species-consistent subclades, including the teleost-specific *Ol* lasp-1A and *Ol* lasp-1B paralogs. Notably, *Dr* lasp-2N was separated from other lasp-2 members, consistent with its lower sequence similarity in both the third nebulin repeat and linker (Fig.3C).

The 3D structural prediction results were consistent with the domain prediction results (Fig. 3E). The LIM and SH3 domains showed similar structures among lasp family proteins. Nebulin repeats formed α-helices with bends around the conserved SDXXYK motif (Fig. 3E). All lasp-1 structures, which contain two nebulin repeats, showed two α-helical regions (Fig. 3E). *Lo* lasp-2, which contains the third nebulin repeat, showed three α-helical regions. *Ol* lasp-2, which has a third nebulin repeat predicted by SMART despite the low homology to tetrapod lasp-2 (Fig. 3A), also formed a corresponding α-helix. *Dr* lasp-2N or lasp-2L, which lack the third nebulin repeat, showed two α-helical regions.

### 2.5 Tissue expression of lasp family proteins and nebulette is similar among zebrafish, medaka, and spotted gar with partial differences

We analyzed RNA-seq data from PhyloFish to examine whether *LASP1* and *LASP2/NEBL* expression patterns were conserved between ray-finned fish and mammals (Fig. 4). *Lo LASP1* mRNA showed ubiquitous expression, except for low expression in muscle, consistent with mammalian *LASP1* expression (Schreiber et al. 1998). However, *Dr LASP1* mRNA showed lower expression in the testis and much lower expression in unfertilized eggs; the level in muscles was similar to that in other tissues. *Ol* lasp-1A also showed ubiquitous expression except for low expression in muscles, similar to *Lo* lasp-1, whereas *Ol* lasp-1B was mainly expressed in the testis, with lower expression in other tissues. Apart from *Ol* lasp-1B, all lasp-1 of these three ray-finned fishes are expressed in the gills, which is fish-specific.

**Fig. 4.**
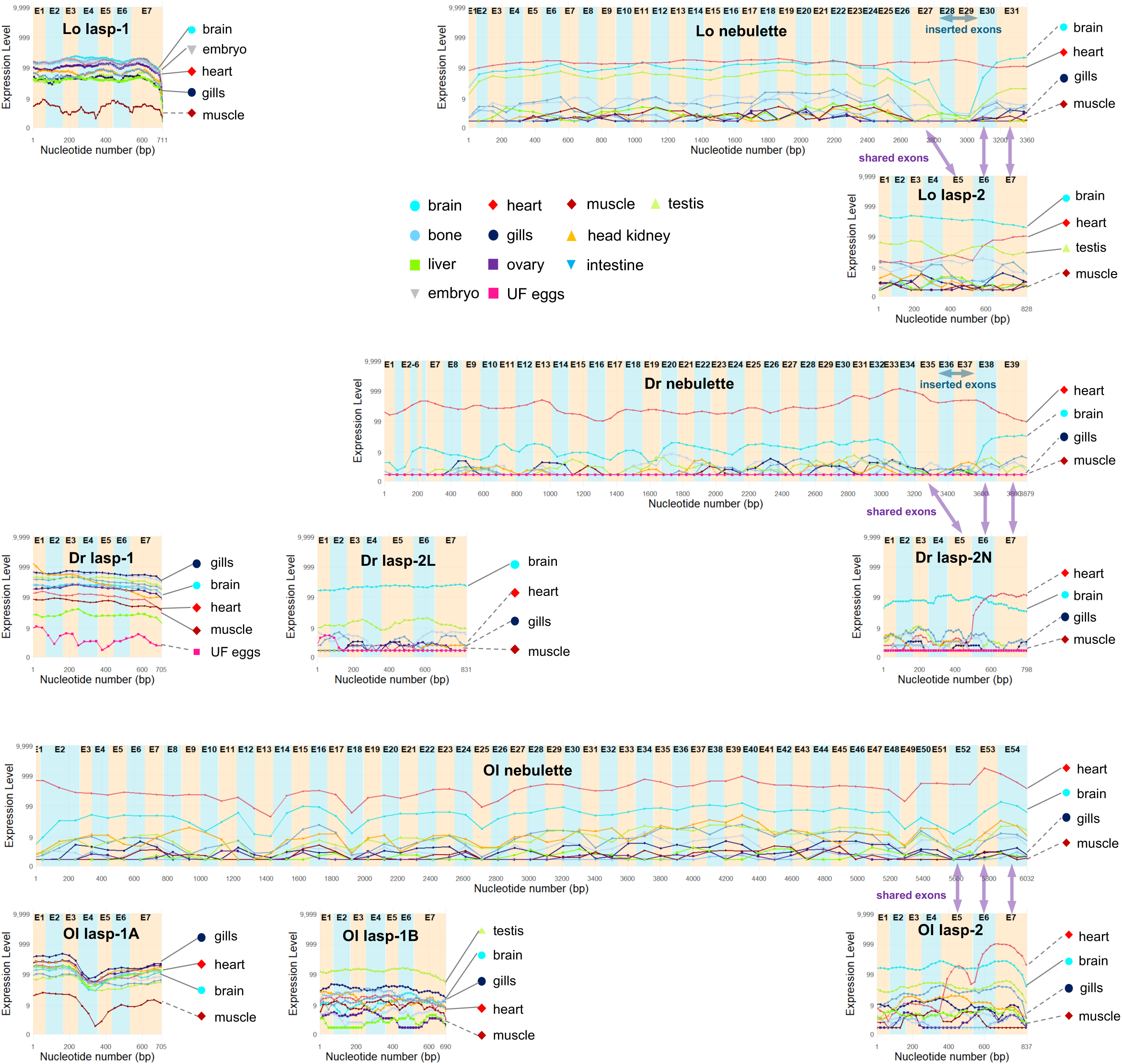
Tissue expression of lasp family proteins of spotted gar, zebrafish and medaka. The horizontal axis represents the position of the coding region of each mRNA and the vertical axis represents logarithmic RPKM of 10 tissues, embryo, and unfertilized eggs (zebrafish). Expression of the brain, heart, and muscle, where expression of nebulin superfamily were reported in humans, is indicated by lines (high/medium expression) or dotted lines (low expression). Additionally, high expression tissues (e.g. embryo, testis, and gills) are indicated in each transcript. “Shared exons” indicate exons 5–7 of *Lo* lasp-2, *Dr* lasp-2N, and *Ol* lasp-2, which are shared with nebulette mRNA. “Inserted exons” of nebulette indicate two additional nebulette-specific exons confirmed in humans, spotted gar, and zebrafish, but not identified in chicken, tropical frog, and medaka.

*Lo* nebulette showed high expression in the heart across the entire transcript (Fig 4), similar to mammalian nebulette (Schreiber et al. 1998). For the brain and testis, *Lo* nebulette showed varying RPKM (number of reads per kilobase per million reads) counts depending on region. We consider that the overall expression is actually low in the brain and testis, because the low overall expression of *Lo* nebulette-specific exons 28 and 29, along with high counts of *Lo* lasp-2-shared exons 30 and 31, may be due to counting of *Lo* lasp-2 mRNA. *Lo* lasp-2 showed the highest expression in the brain similar to that reported in mammals (Li et al. 2004), with medium expression in the heart and testis but lower expression in other tissues. *Dr* nebulette showed high expression in the heart and low expression in other tissues, similar to *Lo* nebulette. *Dr* lasp-2N, a splice variant of *Dr LASP2/NEBL*, showed brain-specific expression. TGD-derived *Dr LASP2L* showed a similar pattern. Comparable to *Lo* and *Dr* nebulette, *Ol* nebulette showed high expression in the heart. *Lo* lasp-2 and *Ol* lasp-2 showed high expression in the brain and moderate expression in the testis.

### 2.6 All zebrafish lasp family proteins have actin-binding activity

Recombinant *Dr* lasp-1, *Dr* lasp-2N, and *Dr* lasp-2L exhibited actin-binding activity (Fig. 5). In the absence of actin, only a small proportion of *Dr* lasp family proteins sedimented, with the majority remaining in the supernatant following high-speed centrifugation. With actin at a 5:1 molar ratio, approximately half of *Dr* lasp-1 and *Dr* lasp-2N co-sedimented with actin, whereas *Dr* lasp-2L showed less co-sedimentation (Fig. 5). Since glutathione S-transferase (GST)-lasp-2N and GST-lasp-2L easily aggregate and become absorbed during preparation, it was difficult to compare their Kd values.

**Fig. 5.**
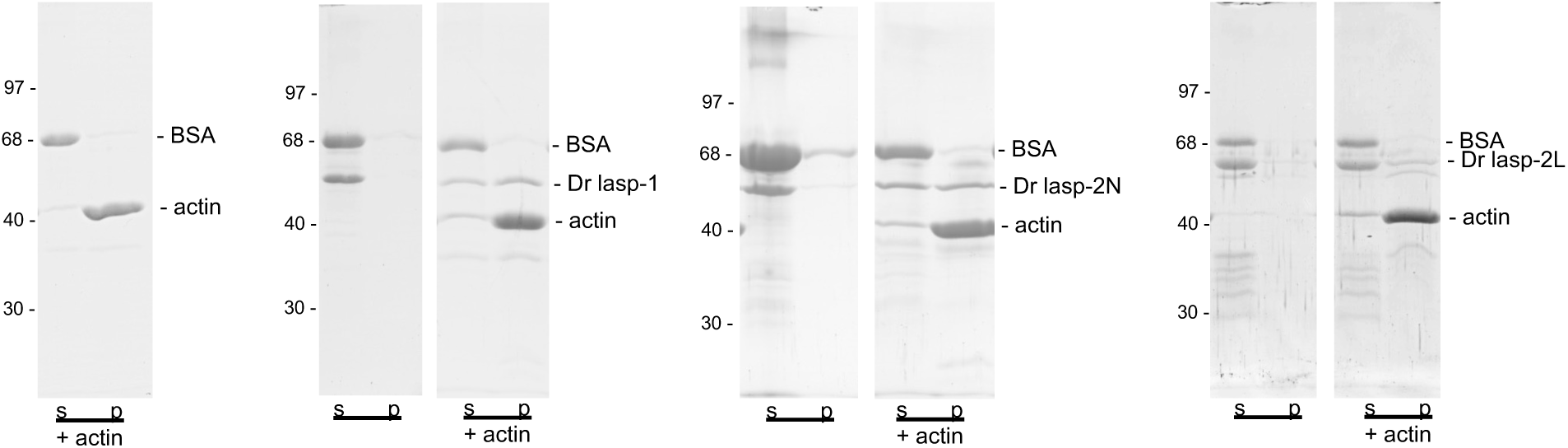
Actin-binding activity of three zebrafish lasp family proteins. Co-sedimentation assay of *Dr* lasp-1, *Dr* lasp-2N, and *Dr* lasp-2L. All lasp family proteins were co-precipitated with actin filaments. Samples were run in a 10% gel and the mobilities of molecular mass markers are listed on the left of the gel images (in kilodaltons). Positions of BSA, zebrafish lasp family proteins, and actin in supernatant (s) and pellet (p) samples are indicated on the right of the gel images. Because BSA was added to GST-*Dr* lasp-2N and GST-*Dr* lasp-2L to prevent adsorption to dialysis and centrifugation tubes, BSA was also added to GST-lasp-1.

### 2.7 Zebrafish lasp family proteins localize in actin-containing structures of cultured cells

We compared the subcellular localization of *Dr* lasp-1, *Dr* lasp-2N, and *Dr* lasp-2L using Lifeact-labeled F-actin to examine whether their localization is conserved between teleosts and tetrapods. We expressed zebrafish lasp family proteins in chicken fibroblasts, a well-established system for studying the actin cytoskeleton. Chicken fibroblasts displayed a variety of shapes, with lamellipodia and filopodia at the leading edge and polygonal cell edges at adhesion sites. All *Dr* lasp proteins co-localized with F-actin in lamellipodia and filopodia (double arrowheads), as well as focal adhesions (open arrowheads) at stress fiber tips, where integrin-mediated cell adhesions between stress fibers and extracellular matrix were formed (Fig. 6).

**Fig. 6.**
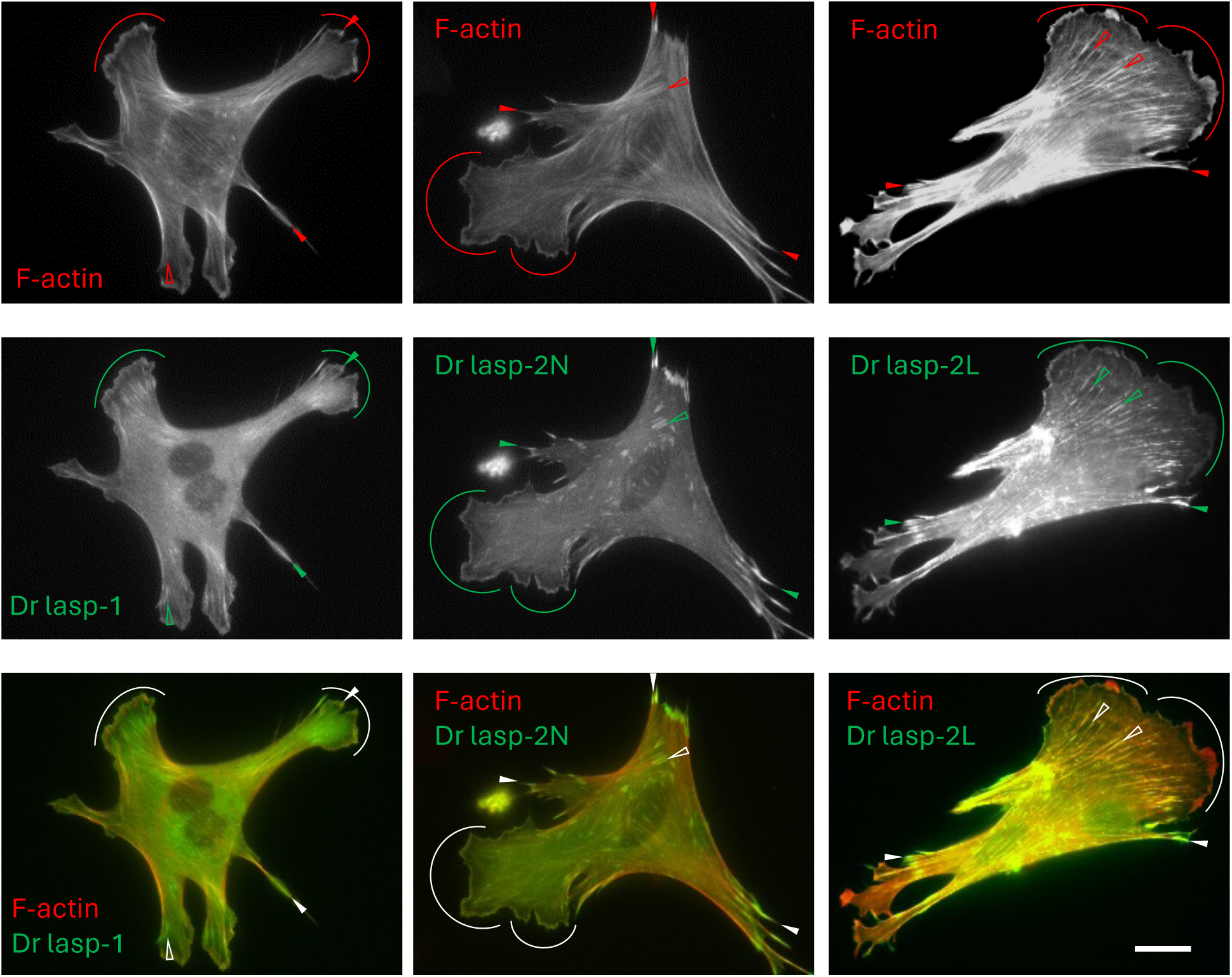
Localization of three lasp family proteins of zebrafish ectopically expressed in chicken fibroblasts. Representative images of EGFP-tagged *Dr* lasp-1, *Dr* lasp-2N, and *Dr* lasp-2L, cotransfected with mCherry-lifeact. Examples of focal adhesions are indicated by blank arrowheads (cell inner) and filled arrowheads (cell periphery). Lamellipodia are indicated by arcs. Bars in all images represent 20 μm.

## 3 Discussion

### 3.1 Evolutionary origin of three lasp family proteins of teleost

In this study, we identified a third zebrafish lasp family gene (*Dr LASP2/NEBL*) on chromosome 24 using the new genome assembly GRCz12tu and PhyloFish, in addition to *Dr LASP1* on chromosome 3 and the gene annotated as “nebulette” in ZFIN on chromosome 2. The gene annotated as “nebulette” lacked nebulette exons and is much smaller than nebulette of other species, so we redefined it as *Dr LASP2L*, encoding lasp-2L (lasp-2-like protein); whereas the newly identified gene (on chromosome 24) generated two mRNA transcripts homologous to mammalian and avian lasp-2 and nebulette (Fig. 1), so we assigned the name *Dr LASP2/NEBL*, encoding lasp-2N (lasp-2-new) and nebulette. We first identified *Dr LASP2/NEBL* in the Phylofish database; however, it was absent in the previous genome assembly (GRCz11) because the gene was mapped near the end of chromosome 24, and GRCz11 lacked this region. Even in the current genome assembly (GRCz12tu), the predicted exon structure differed markedly from the PhyloFish transcripts of Dr lasp-2N and nebulette (Fig. 1). Thus, transcriptome data are important for identifying genes and accurately defining their exon structures.

The three lasp family proteins in zebrafish and medaka are encoded by seven exons, similar to mammals and aves lasp-1 and lasp-2. However, while mammalian and avian lasp-1 and lasp-2 contain two and three nebulin repeats, respectively, all three zebrafish lasp proteins contain two nebulin repeats (Fig. 3A and 3C). DNA-based synteny analysis showed that zebrafish chromosomes 24 and 2, containing *Dr LASP2/NEBL* and *Dr LASP2L*, respectively, were derived through TGD from a genomic region corresponding to spotted gar chromosome 6 (Fig. 2A). This relationship was further supported by gene-based synteny analysis, which showed conserved neighboring genes around *Dr LASP2/NEBL* and *LASP2L* (Fig. 2B). Additionally, the phylogenetic tree of the linker region grouped Dr lasp-2N and Dr lasp-2L with vertebrate lasp-2 (Fig. 3D). Together, these results indicate that *Dr LASP2/NEBL* and *Dr LASP2L* are TGD-derived ohnologs. In contrast, DNA-based synteny analysis readily revealed that medaka chromosomes 8 and 19, containing *Ol LASP1A* and *Ol LASP1B*, respectively, were derived through TGD from a genomic region corresponding to spotted gar chromosome 28 (Fig. 2A). The phylogenetic tree of the linker region also grouped *Ol* lasp-1A and *Ol* lasp-1B with vertebrate lasp-1 (Fig. 3D). Although both zebrafish chromosomes 3 and 12 correspond to TGD-derived regions homologous to spotted gar chromosome 28, only chromosome 3 retains *Dr LASP1*, indicating that the duplicate was lost from chromosome 12 (Fig. 2A). Similarly, although both medaka chromosomes 17 and 20 correspond to TGD-derived regions homologous to spotted gar chromosome 6, only chromosome 20 retains *Ol LASP2/NEBL*, indicating that the duplicate was lost from chromosome 17 (Fig. 2A).

### 3.2 Loss of duplicate genes and exons of zebrafish lasp family proteins

Gene duplication may have driven evolutionary innovation and species diversity. It has been suggested that the duplicate copy either became a pseudogene via mutation or was physically lost (reviewed by Albalat and Cañestro 2016). However, ohnologs may also be retained in two copies and either specialize by partitioning ancestral gene functions (i.e., subfunctionalization) or acquiring a novel function (i.e., neofunctionalization) (Postlethwait et al. 2000). For example, the duplication-degeneration complementation (DDC) model proposes that duplicates can be conserved by complementary degenerative mutations (Force et al. 1999). Divergence of coding regions can be achieved through point mutations, insertions/deletions, exon gain/loss, exonization/pseudoexonization, exon shuffling, and divergence of alternative splicing. Genes usually have multi-functions and the functional unit is commonly exon-coded. If complementary loss of sub-functions occurred, both duplicated genes become fixed in the genome, because loss of either gene would disrupt essential ancestral gene function.

Indeed, in zebrafish, many duplicated genes became pseudogenes or were lost, and exon gain/loss was reported in some ohnologs (Altschmied et al.; Hultman et al. 2007; Rojo-Bartolomé et al. 2020). Also in rainbow trout (*Oncorhynchus mykiss*), around half of the ancestral genes have been retained in duplicates, with the remaining being singletons (reviewed by Glasauer and Neuhauss 2014). This is consistent with our finding that almost all nebulin superfamily genes of zebrafish and medaka exist as singletons except duplicated lasp family genes (Table S1). Additionally, *Dr LASP2L* lacks the nebulette exons, generating a single lasp-2L transcript (Fig. 1); thus, zebrafish have two lasp-2 proteins (*Dr* lasp-2N and *Dr* lasp-2L) and one nebulette protein.

Amino acid sequences of duplicated lasp family proteins are more diverse than singleton lasp. For example, among lasp-1 of ray-finned fish, *Ol* lasp-1B, generated by TGD, has a different linker sequence (Fig. 3C and 3D). Similarly, *Dr* lasp-2N also has a linker sequence phylogenically far from other lasp-2 proteins. These variations may suggest subfunctionalization or neofunctionalization of these TGD-derived lasp family proteins. All ray-finned fish lasp family proteins showed strong conservation in the LIM domain, the first and the second nebulin repeats, and the SH3 domain, suggesting these regions are essential for their functions.

During the exon–intron structural analysis of nebulette in zebrafish and medaka, which shares some exons with lasp-2, we identified several exons predicted in the NCBI genomic annotations that are not represented in the corresponding PhyloFish transcripts (Table S1). Although alternative splicing commonly involves the inclusion or skipping of individual cassette exons, extensive exon skipping can occur in genes encoding mammalian nebulin. For example, Donor et al. (2010) reported skipping of exons 82–105 of mammalian nebulin, coding 4,374 bp (1,458 aa) of nebulin repeats. The length of these skipped exons are an exact multiple of three, called “in-frame exons”, and are suggested to contribute to differences in the length of filamentous structure along the actin filament. Notably, the lengths of the NCBI-predicted but PhyloFish-unrepresented exons of *Dr* nebulette and *Ol* nebulette are all exact multiples of three (data not shown). Additionally, almost all exons coding nebulin repeats in nebulette of ray-finned fish and tetrapods used in our study were in-flame exons (see supplemental data 1); thus, they can constitute splice isoforms without frame-shifts. Their transcripts containing these skipped exons will be investigated in future analyses.

### 3.3 Conservation of actin-binding activity among lasp family proteins

We previously showed that the LIM domain and the first nebulin repeat are the actin-binding region of Gg lasp-2 (Nakagawa 2009). Furthermore, we also confirmed that *Ci* (*C. intestinalis*) lasp and *Ce* (*C. elegans*) lasp, with high sequence similarity in the actin-binding region, exhibit actin-binding activity (Terasaki et al. 2006 ; Fujita et al. 2021). In the present study, as expected from high sequence similarity in the region of the LIM domain and the first nebulin repeat, *Dr* lasp-1, *Dr* lasp-2N, and *Dr* lasp-2L displayed actin-binding activity (Fig. 5). Moreover, some of the phosphorylation sites (S61, S99, S146, and Y171) reported in mammalian lasp-1 (Chew et al. 2002; Keicher et al. 2004;Lin et al. 2004) are conserved in all three lasp family proteins of zebrafish (Fig. 3C), suggesting they are regulated in a similar manner to lasp-1 in vivo. However, the affinity of lasp-2L seemed to be lower than lasp-1 and lasp-2N in the co-sedimentation assay (Fig. 5). The lasp-2L sequence differs from lasp-1 and lasp-2N mainly in the linker region (Fig. 3C), and this may lead to differences in their amorphous 3D structures (Fig. 3E). However, the function of the linker region has not yet been analyzed, even in mammalian and avian lasp-1 and lasp-2. Additionally, the absence of the third nebulin repeat in *Dr* lasp-2N and lasp-2L may also result in differences in actin-binding activity, but to date, the function of the second and third nebulin repeats has also not been analyzed.

The detailed functions of the three zebrafish lasp proteins in actin filaments should be analyzed in the future by observing actin filament structures using immunoelectron microscopy or fluorescently labeled F-actin. For example, our previous studies have used these approaches to analyze the functions of other lasp proteins in detail, reporting weak bundling activity of chicken lasp-2 (Zieseniss et al. 2008).

### 3.4 Universal and specific functions of lasp-1, lasp-2, and nebulette across species

Cellular and biochemical function of lasp family proteins of ray-finned fish have not been analyzed until our current study. However, zebrafish has ZFIN, an excellent integrated database of genes including mRNA expression patterns, gene mutants, and siRNA effects. Among the three zebrafish lasp family proteins, only *Dr* lasp-1 (https://zfin.org/ZDB-GENE-030131-1936) and *Dr* lasp-2L, currently annotated as “nebulette” (https://zfin.org/ZDB-GENE-050320-151), are reported in ZFIN. This, combined with our current achievements in examining tissue expression, actin-binding activity, and localization in ectopically expressed cells (Fig. 5), helps in elucidating functional differences.

In mammals, lasp-1 was reported to be highly expressed in a wide range of non-muscle tissues and is particularly enriched in several actin-rich cell types in mammals (Chew et al. 2000). We found that Lo lasp-1 and *Ol* lasp-1A exhibit a pattern similar to that of mammals, showing high expression levels across various non-muscle tissues (Fig. 4). Additionally, *Dr* lasp-1 exhibits moderate expression in muscle and may regulate myogenesis. Expression *Lo lasp-1*, *Dr lasp-1*, and *Ol lasp-1A* (Fig. 4) in the gills, might be related to the fact that epithelia contain prominent F-actin-based microridges on their apical surfaces (Depasquale 2018; Lu et al. 2022). The low *Dr* lasp-1 expression level in UF eggs was consistent with the low lasp-1 signal in UF eggs following in situ hybridization (https://zfin.org/ZDB-PUB-040907-1#summary), suggesting that lasp-1 expression 1 may regulate embryogenesis (Aanes et al. 2011). Indeed, morpholino-mediated *Dr lasp-1* knockdown resulted in abnormal morphants with defects affecting multiple organs (Grossi et al. 2023). Functional divergence may have occurred between the two medaka *lasp-1* paralogs, as suggested by differences in their amino acid sequences and tissue expression patterns. Prominent testis-specific expression of *Ol* lasp-1B suggests this ohnolog may specifically participate in spermatogenesis, much like lasp is required for spermatid individualization in fruit flies (Lee et al. 2008).

The high expression patterns of *Lo* lasp-2, *Dr* lasp-2L, *Dr* lasp-2N, and *Ol* lasp-2 are largely consistent with those in mammals and chickens (Li et al. 2004; Terasaki et al. 2004). Previous studies analyzing the “nebl” gene annotated in ZFIN provide various information about *Dr* lasp-2L function. For example, low lasp-2L expression in the heart (Shih et al. 2015) and identification of lasp-2L in proteome analysis of postsynaptic density fraction (Bayés et al. 2017) is consistent with our present study confirming that lasp-2L is mainly expressed in the brain (Fig. 4). However, CRISPR knockouts of *Dr* lasp-2L induced prolongation of the ventricular action potential (Maurya et al. 2023); thus, lasp-2L may also regulate cardiomyocyte contraction. Transcriptome analysis showing varied mRNA levels of *Dr* lasp-2L among neuron types (Takeuchi et al. ; Pandey et al. 2018), suggesting that lasp-2L regulates morphology, which differs among neurons. *Dr* lasp-2N and *Dr* lasp-2L may have undergone subfunctionalization, because their amino acid sequences corresponding to the lost nebulin repeat and linker varies (Fig. 3C) and *Dr* lasp-2L seemed to have lower affinity to actin (Fig. 5). However, our newly identified lasp-2N has no information in ZFIN; this is because gene annotation of ZFIN depends on the previous genome assembly (GRCz11tu), which lacks the region where *Dr LASP2/NEBL* is located. *Lo* lasp-2 and *Ol* lasp-2 are also moderately expressed in the testis (Fig. 4). Some studies indicated that neurons and sperm share a number of characteristics, including the importance of the exocytotic process and the presence of similar receptors and signalling pathways (reviewed by Matos et al. 2021). In mammalia and aves, nebulette is a cardiac-specific isoform of *LASP2/NEBL* and localizes to Z-discs of cardiac muscles. Several mutations of nebulette have been identified in cardiomyopathy, suggesting nebulette is an essential component in heart sarcomeres (Perrot et al. 2016). Specific expressions of *Lo*, *Dr*, and *Ol* nebulette in the heart (cardiac muscles) suggest similar functions to those in mammalia and aves (Fig. 4). However, in a similar case to *Dr* lasp-2, our newly identified *Dr* nebulette has no information found in ZFIN. We also should consider that some studies note inconsistencies between mRNA and protein expression (Liu et al. 2016); thus, protein expression in various tissues should be analyzed in the future using specific antibodies.

Three lasp family proteins of zebrafish showed similar localization when ectopically expressed in fibroblasts (Fig. 6), but may show differences in other cell types. In mammalian cell lines, ectopically expressed lasp-1 and lasp-2 showed similar or different localization patterns depending on cell type and cell status (Li et al. 2004 ; Panaviene and Moncman 2007; (Deng et al. 2008)). Additionally, Deng et al. (2008) found that lasp-2 overexpression in PC6 cells (established from neuroblastoma) suppressed neurite outgrowth in response to b-NGF, although overexpression of lasp-1 showed no effect; they also showed that lasp-2 overexpression in NIH3T3 cells (established from fibroblasts) promoted cell spreading, whereas lasp-1 overexpression had no effect. In primary cells, the expression and localization of lasp-1 and lasp-2 differs. For examples, in cardiomyocytes, we previously used specific antibodies to report that lasp-2, but not lasp-1, was localized to the Z-line (Zieseniss et al. 2008), suggesting that the third nebulin repeat or linker of lasp-2 plays specific roles in cell morphology and motility. However, no unique binding proteins have been identified in such specific regions of lasp-1 and lasp-2. Recently, we revealed that lasp-2 accumulated in actin-containing substructures of astrocytes, which are the predominant type of glia in the central nervous system and have long, branched stem processes and perisynaptic/ peripheral astrocyte processes (PAPs) contacting neurons and other glial cells (Ikoma et al. 2026). Furthermore, immunostaining and transfection of lasp-2 in cultured chicken astrocytes showed that endogenous lasp-2 localizes to filopodia and lamellipodia in actively moving PAP-like structures. Additionally, lasp-2 and F-actin colocalized as small elliptical structures at the base of lamellipodia and filopodia of process tips, which may be cell–substrate adhesions.

### 3.5 Diversified functions of actin-binding proteins generated by TGD

Zebrafish is a widely used vertebrate model organism due to established reverse genetics techniques like transgenic, knockdown, and knockout of genes. However, only a few past studies have examined the functional differences among actin-binding proteins generated by TGD. For example, within the zebrafish gelsolin family, the actin-severing proteins gelsolin (*gsna*/*gsnb*), scinderin (*scina*/*scinb*), and the teleost-specific scinderin-like proteins (*scinla*/*scinlb*) are considered ohnologs resulting from TGD. Based on phylogenetic analyses, Jia et al. (2007) showed that zebrafish have at least 6 genes encoding gelsolin-like proteins, and clear functional differentiation is observed among these duplicated genes. Kanungo et al. (2003) showed that inhibition of *gsna* expression resulted in ventralized phenotypes, including the absence of a brain and eyes. They noticed that these phenotypes differed from those of gelsolin-null mice and proposed that multiple gelsolin members may exist. Although *gsnb* suppression results were not reported in the study, *gsnb* tissue expression pattern was found to partially differ from that of *gsna*. Furthermore, the teleost-specific paralog *scinla* has acquired a completely different physiological role, encoding a major crystallin in the cornea (notably, zebrafish also possess a separate gene corresponding to mammalian crystallins). In contrast, *scinlb* lacks corneal-specific expression, suggesting it maintains its original role as an actin-binding and severing protein across a wide range of tissues, including the skin and intestine (Jia et al. 2007).

Such functional subdivision is also evident in other actin-binding families. Within the thymosin β4 family, which binds monomeric actin (G-actin), multiple paralogs (such as *tmsb4x* and *tmsb*) exist in zebrafish. The *tmsb4x* paralog exhibits a highly specialized functional adaptation that promotes actin polymerization and facilitates specific axonal regeneration in the central nervous system (Song et al. 2024). Meanwhile, *tmsb* shows distinct expression domains in the gills and thymus, demonstrating a clear division of labor. Similarly, duplicated genes within the zebrafish α-actinin family exhibit distinct spatial and temporal expression patterns, suggesting functional specialization while retaining the conserved structural and actin-binding properties of vertebrate α-actinins (Holterhoff et al. 2009).

These examples strongly suggest that paralogs generated after TGD acquired distinct physiological roles, contributing to the biological complexity of teleosts. Recently, the updated zebrafish genome assembly, excluding unlocalized scaffolds, was completed (GRCz12tu, released in 2025). Indeed, we could not identify *LASP2/NEBL* in previous assemblies, even though we had identified the lasp-2N and nebulette mRNAs in the PhyloFish database. The updated genome assembly will be critical in the future for determining how many actin-binding proteins remained in teleosts after TGD and clarifying their evolutionary trajectories.

## 4 Materials and Methods

### 4.1 Identification of genes of nebulin superfamily proteins of Teleostei

Gene and transcript sequences of tetrapod *LASP1* and *LASP2/NEBL* were determined in our previous study (Fujita et al., 2022): Mammalia (human, *Homo sapiens* = Hs), Aves (chicken, *Gallus gallus* = Gg), and Amphibia (tropical clawed frog, *Xenopus tropicalis* = Xt). We had also reported the full mRNA sequences of lasp-1, lasp-2, and nebulette of Actinopterygii (spotted gar, *Lepisosteus oculatus* = Lo) from the PhyloFish database (http://phylofish.sigenae.org/index.html) (Pasquier et al. 2016) of tissues and embryos from various fish species. We also reported the gene structures of the transcripts of spotted gar using the NCBI database (fLepOcu1). However, the genome assembly of spotted gar was recently updated to fLepOcu1.hap2 (https://www.ncbi.nlm.nih.gov/datasets/genome/GCF_040954835.1/) in 2024; thus, in the present study, we reanalyzed the sequences and gene structures of lasp-1, lasp-2, and nebulette using fLepOcu1.hap2 (Supplemental Table 1).

Using translated nucleotide BLAST (tBLASTn) and previously identified *Lo* lasp-1 and *Lo* lasp-2 amino acid sequences (Fujita et al. 2021) as queries, lasp family proteins of zebrafish and medaka were searched against two databases : (1) Phylofish database of zebrafish and medaka; and (2) zebrafish genome data of GRCz12tu (https://www.ncbi.nlm.nih.gov/datasets/genome/GCF_049306965.1/) and medaka genome data of Olatipes_Hd-rR_3.1 (https://www.ncbi.nlm.nih.gov/datasets/genome/GCF_053564925.1/). Additionally, NRAP and nebulin of spotted gar, zebrafish, and medaka were searched against the above genome and transcriptome databases using BLASTn, with human N-RAP and nebulin (see Table 1) as queries. Based on the domain definitions provided by the SMART program (described in Section 5.2), transcripts encoding proteins equipped with a complete set of a LIM domain, nebulin repeats, and an SH3 domain were annotated as lasp family proteins. Furthermore, transcripts whose protein products share the same SH3 domain as lasp-2 were identified as nebulette. Transcripts encoding proteins that possess only a LIM domain and multiple nebulin repeats (>4) were designated as N-RAP, whereas those containing multiple nebulin repeats (>4) and an SH3 domain were classified as nebulin. We also searched ZFIN using keywords “neb/nebulin”, “nrap/N-RAP”, or “nebulette/nebl”.

The NCBI genome data for zebrafish (Tuebingen strain) and medaka (Hd-rR) are highly reliable because they are derived from inbred strains and lack unplaced scaffolds. For the spotted gar, the genome assembly contains 784 unplaced scaffolds; while future re-analyses may shift global chromosomal positions, the local synteny of genes within the sequenced regions is unlikely to be affected. Although RefSeq gene annotations are complete for all three species, computational exon predictions may sometimes be inaccurate, as observed with *Dr LASP2/NEBL*. In contrast, because the PhyloFish database is based on SRA data, it represents actual transcribed mRNAs. Thus, this database accurately reflects true exon structures and genetic polymorphisms (such as SNPs) rather than computational predictions. Therefore, in our study, the PhyloFish database was prioritized for determining actual exon-intron boundaries. However, when partial sequence discrepancies occurred between the PhyloFish and NCBI databases, NCBI genomic sequences were prioritized for determining the final amino acid sequences.

### 4.2 Analysis of genomic organization, domain, and 3D structure

The exon numbers, intron lengths, and genomic sizes of lasp family proteins in zebrafish spotted gar and medaka were analyzed by comparing mRNA and genomic sequences, and intron positions were determined manually according to the GT-AG intron rule.

Amino acid sequences deduced from transcripts predicted in the genome database and the phylofish database were analyzed using the InterProScan platform at the European Bioinformatics Institute (http://www.ebi.ac.uk/Tools/pfa/iprscan/) according to the definitions of the SMART domain identification program (proteins with a LIM domain, nebulin repeats, and SH3 domain).

Using AlphaFold v3, a program implemented in a web-based server AlphaFold Server (https://alphafoldserver.com/welcome), we analyzed the 3D structures of lasp proteins of *Lo* lasp-1, *Lo* lasp-2, *Dr* lasp-1, *Dr* lasp-2N, *Dr* lasp-2L, *Ol* lasp-1A, *Ol* lasp-1B and *Ol* lasp-2 proteins using default settings.

The 3D structures of the lasp family proteins, predicted via AlphaFold, were visualized and optimized using the PyMOL Molecular Graphics System (Schrödinger, LLC). Specific structural features, including the LIM domain, nebulin repeats, and SH3 domain, were highlighted using custom color palettes, with the remaining protein backbone colored light gray. All protein models were displayed in a cartoon representation. For accurate size comparison across different structural models, a 50 Å scale bar was computationally generated within the 3D coordinate space using pseudoatoms. Finally, the camera was set to orthoscopic projection to eliminate perspective distortion, and high-resolution images were rendered utilizing PyMOL’s built-in ray-tracing algorithm with a transparent background.

### 4.3 Synteny analysis

We analyzed BLAST-based orthology between spotted gar and zebrafish using both DNA and amino acid sequences. For synteny analysis based on DNA sequences, we used the Comparative Genome Viewer (https://www.ncbi.nlm.nih.gov/cgv/), a visualization tool for whole-genome assembly-assembly alignments provided by NCBI, to compare chromosome 6 or chromosome 28 of spotted gar (fLepOcu1.hap2) with whole chromosomes of zebrafish (GRCz12tu), setting a minimum alignment length of 10,000 bp.

For synteny analysis based on amino acid sequences, we developed a biopython-based program and ran it on Google Colab. We used spotted gar chromosome 6 (NC_090701.1), where *Lo LASP2/NEBL* is located, and spotted gar chromosome 28 (NC_090723.1), where *Lo LASP1* is located, as query sequences. Using the query sequences, we searched for homologous genes in three zebrafish chromosomes of interest: chromosome 3 (NC_133178.1) containing *Dr LASP1*, chromosome 24 (NC_133199.1) containing *Dr LASP2/NEBL*, and chromosome 2 (NC_133177.1) containing *Dr LASP2L*. Briefly, amino acid sequences in NCBI RefSeq records for protein-coding mRNA transcripts (accession numbers beginning with the prefix “NP”), which have been experimentally analyzed, as well as model RefSeq records (accession numbers beginning with the prefix “XP”), which are predicted in the NCBI annotation pipeline, were extracted from whole sequences of each chromosome. Then, using amino acid sequences of all genes on spotted gar chromosome 6 or 28 as queries, we conducted BLASTP searches against amino acid sequences (derived from genes with accession NM or XM) from the three zebrafish chromosomes of interest, followed by filtering with an E-value threshold of 1e-15 and a minimum coverage of 50%. Most searches yielded single-hit matches. In some cases with multiple hits, a single homologous gene was selected by first evaluating the lowest E-value, followed by the highest bit score, and finally the highest % identity. Then, we confirmed the expression of each homologous gene using a BLASTN search against mRNA sequences from Phylofish. We mapped their chromosomal locations around the three zebrafish chromosomes of interest.

### 4.4 Alignment and phylogenetic tree analyses

Multiple sequence alignment was performed using ClustalW in MEGA X (https://www.megasoftware.net/) and re-aligned manually. Datasets of the third nebulin repeat and linker sequences were analyzed separately using the maximum likelihood (ML) method with bootstrap values (percentage of 100 replicates) displayed in MEGA X. The LG model was used as the amino acid substitution model. Evolutionary rates among sites were modeled using a gamma distribution (G) with five discrete gamma categories. Positions containing gaps or missing data were treated using partial deletion with a site coverage cutoff of 70%. The Nearest-Neighbor-Interchange (NNI) algorithm was employed for the ML heuristic search, with the initial tree generated automatically by MEGA X using the NJ/BioNJ algorithms.

### 4.5 Analysis of Tissue Expression

From the “Depth View” function of the PhyloFish database, RPKM values of transcripts of lasp family proteins and nebulette in 10 tissues (ovary, testis, brain, gills, heart, muscle, liver, head kidney, bones, and intestine) and embryo of spotted gar, zebrafish, and medaka were downloaded. Zebrafish possesses an additional dataset for unfertilized (UF) eggs. To ensure sequence accuracy, a blastn search was performed against the respective genomic sequences, and only transcript regions that exactly matched the genome were extracted for subsequent analyses. The sequences corresponding to coding regions were selected and rearranged into graphs, with the horizontal axis representing the number of mRNA bases and the vertical axis representing a logarithmic representation of RPKM using R (https://www.r-project.org/).

### 4.6 Expression vectors of *Dr* lasp-1, *Dr* lasp-2L, and *Dr* lasp-2N

EST clones of *Dr* lasp-1 (IMAGE:6959759, https://www.ncbi.nlm.nih.gov/nuccore/BC066574) and *Dr* lasp-2L (IMAGE :6898261, https://www.ncbi.nlm.nih.gov/nuccore/BC090509) were purchased from Openbiosystems (http://www.openbiosystems.com/). The cDNA of full-length *Dr* lasp-1 and *Dr* lasp-2L were amplified by PCR from the EST clones using the following primer sets containing BamHI and EcoRI sites: (5′-CGCGGATCCAAAATGAACCCGCTGTG-3′ and 5′-CCGGAATTCTTACATGGCCTCCACGTAG-3′) and (5′-CGCGGATCCAAAATGAACCCTCATTGCG-3′ and 5′-CCGGAATTCGAGCGAATGCCCTCAACATAG-3′). Full-length cDNA clone of *Dr* lasp-2N was obtained by RT-PCR amplification from adult zebrafish oligodT-primed brain cDNA, using the following primer set (5’-CGCGGATCCGAGATGAATCCTCCG-3’ and 5’-CCGGAATTCGCGGCTTAGTTCACAC-3’). The PCR products were subcloned into the EcoRV site of the pBluescript vector, and the sequences were confirmed.

For recombinant protein preparation, *Dr* lasp-1, *Dr* lasp-2N, and *Dr* lasp-2L cDNAs cloned into pBlueScript vectors were subcloned into pGEX6p expression vectors (Amersham Biosciences). For analysis of cellular localization, BamHI fragments of pBluescript SK (+) vector were inserted into the BamHI sites of pEGFP and pmCherry vectors. pmCherry-lifeact was described previously (Ikoma et al., 2006). Plasmids were purified using NucleoBond Xtra Midi (Macherry-Nagel) according to the manufacturer’s protocol.

### 4.7 Purification of recombinant proteins

pGEX6p expression vectors of *Dr LASP1*, *Dr LASP2N*, and *Dr LASP2L* were transfected into BL21. Recombinant proteins fused to GST were expressed in bacteria and purified using Glutathione-Sepharose 4B columns (GE Healthcare, Waukesha, WI) according to the manufacturer’s instructions. Recombinant proteins were dialyzed against F-actin buffer (20 mM Na-PO4 and 0.1 M KCl, pH 7.2). Bovine serum albumin (0.2 mg/mL) was added just after elution of *Dr* lasp-2N and *Dr* lasp-2L because these proteins were easily absorbed into the dialysis tubes and centrifugation tubes.

### 4.8 Actin-binding assay

The actin-binding activity of the recombinant proteins was analyzed as previously described (Nakagawa et al., 2009). Briefly, proteins were dialyzed against F-actin buffer (20 mM Na-PO4 and 0.1 M KCl, pH 7.2). After centrifugation at 100,000 × g for 1 hr, each of the supernatants was mixed with 0.2 mg/mL F-actin in polymerization buffer containing 5 mM MgCl2 and 0.2 mM ATP. After incubation for 2 hr at room temperature or overnight at 4℃, the mixtures were centrifuged at 100,000 × g for 1 hr, and the supernatants and pellets were subsequently analyzed using SDS-PAGE. Since *Dr* lasp-2L and *Dr* lasp-2N were dialyzed and stored with BSA, we also added BSA to *Dr* lasp-1 to ensure consistency in subsequent comparisons/analyses.

### 4.9 Expression of zebrafish lasp family proteins in chicken fibroblasts

Primary cultured fibroblasts were prepared from embryonic chick skin as previously described (Murk et al. 2009), with slight modifications. Fertilized eggs of White Leghorn chickens were incubated for 10–11 days with intermittent rocking at 38°C. After removing the embryos, isolated skin pieces were rinsed with phosphate-buffered saline, dissected, treated with 0.25% trypsin-EDTA (Wako, Osaka, Japan), and dissociated by pipetting. Enzymatic dissociation was inactivated by adding Dulbecco’s modified Eagle’s medium (DMEM) containing 10% fetal bovine serum (FBS). Cells in the suspension were dispersed by pipetting several times, centrifuged at 400 ×g for 5 min, resuspended in DMEM containing 10% FBS, 1% chicken embryo extract, 100 IU/mL penicillin, and 100 μg/mL streptomycin, and plated on culture dishes (IWAKI, Tokyo, Japan) coated with 0.5% collagen (IPC-50, KOKEN, Tokyo, Japan) at a cell density of 2.0–4.0 × 104 cells/cm2. The cells were then incubated at 37°C in 5% CO2.

For transfection experiments, freshly prepared fibroblasts were harvested after 1–3 days of culture and treated with 0.25% trypsin-EDTA (Wako, Tokyo, Japan) for 3 min. DMEM supplemented with 10% FBS was added, and the cells were dissociated by pipetting. The dissociated fibroblast cells were collected by centrifugation and transfected using an Ingenio Electroporation Kit (Mirus Bio, Madison, WI, USA) and a Nucleofector device (LONZA, Basel, Switzerland) with the A-24 program, according to the manufacturer’s instructions. The transfected cells were plated on 12 mm glass dishes coated with collagen at a cell density of 1.0–4.0 × 104 cells/cm2 and observed after two days.

## 4. 10 Fluorescence microscopy

Transfected cells were observed using an inverted fluorescence microscope (Axiovert 135, Carl Zeiss, Oberkochen, Germany) equipped with a 40× phase-contrast objective lens, fluorescence filters (Zeiss 39, 38HE, and 43HE), and a CoolSnap HQ CCD camera (Teledyne Photometrics, Tucson, AZ, USA) operated with IPLab software (Scanalytics, Billerica, MA, USA), or a Retiga R1 CCD camera (Teledyne Photometrics) operated with the micromanager in ImageJ software (NIH, Bethesda, MD, USA).

---

**Table S1.** mRNA sequences of nebulin superfamily proteins used in this study. Multiple TSA clones correspond to nebulin of zebrafish and medaka, and we selected the longest TSA clone that has both nebulin repeats and the SH3 domain. The number of nebulin repeats of the curated RefSeq sequence (NM) was cited from NCBI, and the number of nebulin repeats of predicted RefSeq (XM) and TSA were determined using SMART.

**Table S2.** Genes used in the syntenic analysis. Bold letters indicate genes located within a 2-Mb distance from genes of lasp family proteins. We confirmed that all genes used for synteny analysis were expressed using the Phylofish database. Only IRRC3CB close to *Dr LASP1* has no Phylofish TSA clone but mRNA was experimentally obtained.

**Data S1**. Amino acid sequences used in our study.

## Acknowledgments

We thank Dr Yoshinobu Uno for the discussions and suggestions on analysis of zebrafish genome assembly.

## Author contributions

Conceptualization: Y.X., K. M., R.T., Y. F, C. T., A.G.T.; Methodology: Y.X., K. M., R.T., A.G.T.; Formal analysis: Y.X., K. M., R.T., A.N., A.G.T.; Investigation: Y.X., K. M., R.T., Y. M, H. A., S. H, Y. F, C. T., Y.U., Y. K, A.G.T.; Resources: A.N., Y. U., Y. K.,.; Data curation: Y.X., K. M., A. N; Writing - original draft: Y.X., R.T., A.G.T.; Writing - review & editing: Y.U., Y. K, A.G.T.; Visualization: Y.X., H. A., A.G.T; Supervision: Y.U., Y. K, A.G.T.; Project administration: A.G.T.; Funding acquisition: A.G.T.

## Funding

This work was supported by JSPS KAKENHI grant number 22570072, Initiative for Realizing Diversity in the Research Environment of the Chiba University, and Myohara Maroko Research Encouragement Grant.

Academic English Consultation in Chiba University supported English editing.

