## Supplement Data 1 for "Lasp family proteins generated by teleost-specific genome duplication"

**Supplemental Data SD1: Amino acid sequences used in our study**

Exons are represented using letters with/without underlines alternatively.

Double underline: amino acid encoded by two exons.

**Bold letters**: exons shared between lasp-2 and nebulette.

Highlighted letters: amino acid sequences of LIM domain, nebulin repeat (connecting nebulin repeat), and SH3 domain analyzed using the InterProScan platform according to the definitions of the SMART domain identification program.

**>Dr lasp1** 234aa

MNPLCSRCNRVVYPTEKVNCLDKYWHKGCFSCEVCKMTLNMKNYKGFEKRPYCNAHYPKTSFTSVADTPENLRLKQQSKMQSQVLYKEEFEKNKGKGFSVVADTPELQRIKKTQDQISNIKYHEDFEKSRSGGDTPLPLTPQLNTPPAYPSSAASQNYHYEPEPVRPAAAAPPPSSGKRYRAVYDYTAADEDEVSFMDGDMIVDVQQIDEGWMYGRVERTGQQGMLPANYVEAM

**>Dr lasp2L** 276aa

MNPHCARCGKIVYATEKVNCLDKYWHKGCFHCEVCKMTLNMNNYKGYEKKPYCNAHYPKQTFTIVADTPENLRLRQQSELQSQVKYRKDFEQSKGRGLKFVLDTPELQRLRKAQDQISNAKYHKDFEVSGLQGVTQGMRGTYAVRSQMTPNNETVANSTHKAVHQYIMEMDRRPGVIVAPVLPGAYYPNSHSHGHSYMHQTSMHSLRSMQLRSQQMSMTVYRALYDYMAQDYDEVSFRDGDIIQNVQPIDEGWMYGTVQRTGRSGMLPANYVEGIR

**>Dr lasp2N** 265aa

MNPPCARCGKTAYPTEKISCLDKNWHKGCFHCEVCKMTLNMKNYKGYDKKPYCSAHYPKTSFTIVTDTPENLRLKQQSELQSQVKYKRDFEESKGRGFSIVTDTPELERLRKTQEHISN**AKYHEEFERTRGRGATHTLDEQHMEQMSGGSHRGEMERRSAGAAGVFQHHHHLLHHQAPPPQQLLQQGYGQAPAVRSLQSPPASASMQRVYRALYDYSAQDHDEVSFRDGDVIINAQRIDDGWMFGTVQRTGRSGMLPANYVECVN**

**>Dr nebulette** 1292aa

MSDPADASGHTDGGAGAAAESARAADTDPTPDLTSVTSELEVADSQTDAAEENTESQTGGAEEHTEVEDANAQTELETTDALIKKKESSSSRQKRLSSVCVAETLEIEHAKAASQLYSELQYREDGRKDLCVNLYSIMPDTRDTEFAREMTEQQSEVSYRADGRRKLSSSLYSQLPDTPDTQHAKHMTELQSHNKYTADGRKAASVCVFGQLAETLETKHAKEANQLQSQLLYKQAQAGSLYSALPETPEIQLAREITESISENKYSEQSRKSISSCVYSQLPETAETQFSRSISDMQSQVKYKADAQQQMSRSLYHQLSETPETQHARDAAHLQSQVKYKQKVNASSLYSRLPETPETRLAREQTDACSSVRYKAAGRAAAGVSAYSQLAQTQDTLMAKEQSALQSQYEYRRESQQDQCVCVYSQLPETLDTQFVKELTPLISENKYKEEGKKEMQKSLYSSLPHTRETQHARELTQLYSQKAYKEVCKQQAQACVYAQMPQTIETVFAKEVSKQQSDKLYKEKFNSEKGKSSYTQMKDLPDVTHAIQINKQQSDVAYRRGKDELHRYSTVTDRPDILNATHASRLASDVAYRSKADTFDQHTLLQRPDIQHARDAAKLASQVNYKQDLKGQKPQYNPSECVSFRHTQAASALASQVKYSQRKLEAVTDLPNLLQLGHALHASKLQSNVEYRKKYEESKGRYHLALDTAEQRHHKENATLHSQVKYKEEYEKSRGKTQLEFVDTQSYRVSKEAQKMQSEKQYRQQYERELKGSLHLEAELTPMFLNARNASSLISQKEYRKDLEQEVIGKGLMSLEETPEMLRVKNAGHILSEKKYRKDLECEIRGKGADVTSDSLEIQRAKRASEISSDVSYRQASHTGSSFGTVTDTPALLHAAYLRDVYSQKKYRSDVERLKGSCSAASHTPEMERVKNNQRNISALFYTWDSRLMKGLMSTVSDTPEIRLARENAKNISNVQYRDSVGSGTAVADTPEMERVRRNQTNISSVKYKAPVDPVSVGVTPEIERVKQNQQNISTVQYQRGLQEVKGHSCSELDTPELRRVRKSQDSVSM**AKYHEEFERTRGRGATHTLDEQHMEQMSGGSHRGEMERRSAGAAE**LRVWRTDPGSIFDFDPLEDNIQSKSLRRMSERVSQLSSADQSVCSSALWDHQCVERSSVSACSSVCVLQDQPSA**GVFQHHHHLLHHQAPPPQQLLQQGYGQAPAVRSLQSPPASASMQRVYRALYDYSAQDHDEVSFRDGDVIINAQRIDDGWMFGTVQRTGRSGMLPANYVECVN**

**>Ol lasp-1A** 234aa

MNPLCSRCNLVVYPTEKVNCLDKYWHKGCFSCEVCKMTLNMKNYKGFEKRPYCNAHYPKTNFTCVADTPENLRLKQQSKMQSQVLYREDFEKNKGKGFSVVADTPELQRIKKTQDQISNIKYHEEFEKRKMGGEAPLTPTTPSAGAFQPPATPQNYHYEPTPEPVRQAAAAPPPSSGKVYRAVYDYSAADDDEVSFMDGDVIEDVQQIDEGWMYGRVQRTGQQGMLPANYVEAI

**>Ol lasp-1B** 229aa

MMNPSCGRCNKPVYPTEKINCLDKFWHKCCFSCEVCKMALSMKNYKGFEKRPYCSMHYPKTSFTIVTDTPENLRLKQQSQLNSQALYKEDFEKNKGKGFSVVADTPEMQRLKKTQDQISNIKYHEEFEKSKSRSDAPPPENRKDLEKQQNPQRSVQPAAEMQPAAIAPPSGGKRYQALYSYTAAESDEVSFQEGDLIFDAEPIDEGWMYGSNQRTGQRGMLPLNYVRPL

**>Ol lasp-2** 278aa

MNPQCARCGKIVYPTEKVSCLDKNWHKGCFHCEVCKMTLNMKNYKGYDKKPYCNAHYPKTSFTIVADTPENLRLRQQSELQSQVKYKKDFEESKGRGFSIVSDTPEMQRLRKTQEQISN**AKYHEDFERARGRGYTPGLDDPSMERYQRANQMIGEAGYSKGLHSQGMEMDRRPGGIIVAPVLPGAYHHQGAQHQQYHGYMHQTSMSSVRSVTSPPQHSHVRVYRALYDYTAQDHDEVSFRDGDVIINAQPIDEGWMYGTVQRTGKSGMLPANYVESCN**

**>Ol nebulette** 2010aa

MERQAEVEDVKEEEPPAAVEEDEPKSSGDMTETSLASVGSEEEVMEDEGEKKSREEMDNDESEIEKGEEEEHADEETKAKKLMEAWKSEASADAKPPLPDPAFNRRCSLIASDVRYKEDFEKMKGQSLFVPGAELIHSKNISAVISESKYKEEGKKEASLSLYSILPETPETQHAREASELQSEIKYKGNTKTEISSSLYSLLPETTETQFVKELTEMLSENKYKEEGKREMSSSLYSQLPETSEMQLAKTVREFQSEKKYTEDGRRKASISLYSQLPDTPELQHALEVSQLQSQVNYRPKLMRKGTPSSLYSQLPDTTEIQFAREMTELHSESKYKEGGRRSLSQSFYSQLPETAETQFARAVAELQSEMRYKESGRKGVNSSLYSVLPETIETARAKEVSELCSELKYKEDGKKEMNISLYSLLPETMETQHAKEVSDLQSEVKYKDGLKQRIQHSLFHQLPQTTETHLAKQLSELQSQNKYKEVGKKEVKTSLYSLLPQTMETQRAKEASELLSEIKYKESGRKDLSSSLYHTLPDTLETAFAREVTDMLSESKYKDGKKNLAQSFYSQLSQTAETQFAKTVSELQSEVKYKEAGRKGAMSSMYSTLPETLETKHAKDASQLQSQLKYKQSLATEGPSLYHSMPETFDTQFAREQMEMLSEVKYKEDGKKDLSVSLYSVLPETLETQRAKELTEMLSEAKYKEESKKQSSAPLYHLLPETLETRRAKEAALLQSQVLYKEAVKKELSCPVYHKLPETLETQFAQEVTDLQSQNKYKEAGMQSLSHSVYSQLPETEEIKFAKAVADLQSELKYKKWGRQDAGSCLYSEMPETLETQHAKQVSQMQSQVKYKEDVQKQQQSSLFSSLRQTPQTELAKEAAALQSEVKYKGGCKKDMSSPLYHHLPETAETQFAKQMSEFQSELKYKKDMEDLPDTLYSVLPETLETQFVKEMSETHSQVKYKEAVKKEGSSSLYHQLPETLETQRVKEVTELQSENKYKQSGKESMSSSLYAQLPETAQTQLAAKMSELQSESKYKADGLKCLPQSLYSHLPETNETQFVRSVSELLSEAKYKEASKKEESSCLYHQLPETLETLHAKEATELQSQVKYKEGKKDLSSSLYHLLPETEETKFAKTVTELQSENKYKEKCRQEVGSGVFHQMPETKEMEFVKQISELQSESKYKKDVKDLPNSLYSQLPETLQTQFIKEMADKHSQTKYKEASRKELVKSLYSLLPETKETQHAKEQSQLSSERAYREEGKREAGSSLYAQMPQTIETVFAKELTKAQSDKYYKEKYNREKGKADYTNMKTLPDVQHAMEVTKNQSEVGYRRGKEELHHCNMVPDRPDMLNAANAAKLASDVAYKNHTKLPVYSDCSLLTRTDIQHANEVSKLTSQVKYKENFDRHLKGQRPSYNPLDCLSFRHTQAAGALASQVKYRTNQNQRPEGSLDLQLEHALHASKLQSNAEYKKKYEQTKAYYHMAVDTAEQLHHKETAVLHSQVKYREEYEKNRGRSQMEFGETPAYKVSKEVQKMQSEKEYRRDYEEHVKGKALVEVDQTPGYLTARHAGSLLSEKEYKKDLEQEVKGRGLSGVGLEDTPELLRVKKANRILNQKEYRKDLERDILGKGMELSADVLEMQRAKRASEIQSQRSYKQMEQLRENSYGTLTDTPELLHASYLRDIYSQKKYKDDAERLNERYSLLSETPEMERVRANQRHFSSLRYSWDSKLMRGLTTSVTDTPEIVLARENAKKISNVQYREEVGLGTAVTETFEMERVRRNQENISSVKYKQSPAKATSVGVTPEVERVKQNQENISSVKYQRGLQELRGRSCSELDTPEYRRVRKTQDSVSM**AKYHEDFERARGRGYTPGLDDPSMERYQRANQMIGEAGYSKGLHSQGMEMDRRPGGIIVAPVLPGAYHHQGAQHQQYHGYMHQTSMSSVRSVTSPPQHSHVRVYRALYDYTAQDHDEVSFRDGDVIINAQPIDEGWMYGTVQRTGKSGMLPANYVESCN**

**>updated Lo lasp-1** 236aa

MNPSCSRCNKVVYPTEKVNCLDKVWHKACFSCEVCKMTLNMKNYKGFEKKPYCNAHYPKTTFTSVADTPENLRLKQQSKMQSQVLYKEEFEKNKGKGFSVVADTPELQRIKKTQDQISNIKYHEDFEKSRMGGEGHSQTPPPHGTPGYHQQPGAPQNFNFEPAPEPPRQAAAAPPSSAGKRYRAVYDYVAADEDEVSFQDGDMILDVQQIDEGWMYGRVERTGQQGMLPANYVEAI

The sequences below are the same as the supplemental data of our previous study (Fujita et al., 2022).

**>Lo lasp-2** 275aa

MNPQCARCGKIVYPTEKVNCLDKYWHKGCFHCEVCKMTLNMKNYKGYEKKPYCNAHYPKQSFTIVADTPENLRLKQQSELQSQVKYKKDFEESKGRGFSIVSDTPELQRLRRTQEQISN**AKYHEDFEKTRGKGFTPVMDDPTMERARRNHQMVSDAAYKGVQPHVVEMDRKPGIIVAPVLPGAYHPGGHSQGYKYMHQTSVSSMKSMQFPPHSAPMQRVYRAIYDYAAQDHDEVSFRDGDVIINVQPIDEGWMYGTVQRTGKSGMLPANYIETFN**

**>Lo nebulette** 1127aa

MKLCDSKDMSEEDTGVNMKKEEAFSKPIVEDRNLEFAKKCSELISDVKYKEEYEKSKGKCCFVPDTPEFQHVKNLGAFISESKYKEAGKKDISSCLYSQMPETNETQHAKELSQILSEAKYKENGKREFSNCLYANLPETSEIQFAKELSEIQSERKYKKDVKREFSSCLYAKMSNTTETQFAKEMSENLSESKYKQAGKREMSTCLYAQMRETTETQHAKELSQILSEKNYKESGKKDIPFCLYAQMPQTIETVFAKEVAHNQSKILYKQKYDAEKGKSDYSNMKEPPDVAHAMEVNKHQSNVTYKKGMQDIHKYKDVADRLDIRNATETTKLISDVAYKSKMKADVYNHQSVLGRSDIEHAKEASKLVSQVKYKEKFERDLKEKRHQYNPGESVSFKQTQAATALASQVKYKTGLQQMHEPSTDLPNTLHLEHALQATKLQSAIEYKKQYEESKGHYHIALDTAEQLHHKENAVLHSQVKYKEDYEKSKGKSMLEFVDTQTYQVSKEAQKIQSEKEYRREYVDNIKGKAMVDLELTPGYLHARHASTLLSEKEYKKDLETSVKGKGLTGLEETPDFIRVKTAGQILSEKEYRKDLENEIKGKGMEVSTEIPDIQRAKKASQMASQKEYRKDLETEIKGKGMELSTDVLEIKRVKRASEIASQASCKHQLRENMYGTVTDTPEMLHAAYLKDMYSQKKYKHEAEKLKSSYAVVAETPEMQRVKTSQKNVSSLQYNKDKMLLKGTLSSVAETPEILLAKENSKRISNVCYKEGVGSGTAVKDTPEIERVKKNQENISTVKYKAGVGQATAIPDPPELKRVKENQRILSNVNYKAPMGRATAVEMTPEMERVKKNQENISSVKYHSDLKELKGTACAVTDTPELRRVRKTQSDISM**AKYHEDFEKTRGKGFTPVMDDPTMERARRNHQMVSDAAYKGVQPHVVEMDRKPGIIV**DLKVWRTDPGSIFDFDPVEDNIQSKSLHKLSEKANRINRQRSQSVSTVEDDRSELSEKLLSFSGYSTSTARASEDR**APVLPGAYHPGGHSQGYKYMHQTSVSSMKSMQFPPHSAPMQRVYRAIYDYAAQDHDEVSFRDGDVIINVQPIDEGWMYGTVQRTGKSGMLPANYIETFN**

**>Gg lasp-1** 258aa

MNPNCARCGKIVYPTEKVNCLDKFWHKACFHCETCKMTLNMKNYKGYEKKPYCNAHYPKQSFTMVADTPENLRLKQQSELQSQIRYKEEFEKNKGKGFSVVADTPELQRIKKTQDQISNIKYHEEFERSRMGPSPSEGAEMERRNSQESANYRRPEQHQPQHHVQPSNPVYQQQPPASQSYGYKEPAAPASTQRSAPSAAGKRFRAVYDYNAADEDEVSFKDGDTIINVQPIDDGWMYGTVQRTGKTGMLPANYVEAI

**>Gg lasp-2** 273aa

MNPQCARCGKVVYPTEKVNCLDKYWHKGCFHCEVCKMALNMNNYKGYEKKPYCNAHYPKQSFTTVADTPENLRLKQQSELQSQVKYKRDFEESKGRGFSIVTDTPELQRLKRTQEQISN**VKYHEDFEKTKGRGFTPVVDDPITERVRKNTQVVSDAAYKGVHPHIVEMDRRPGIIVAPVLPGAYQQSQSQGYGYMHQTSMSSMRSVHSQPHPAGLRTYRAMYDYSAQDEDEVSFRDGDYIINVQPIDDGWMYGTVQRTGKTGMLPANYIEFVN**

**>Hs lasp-1** 261aa

MNPNCARCGKIVYPTEKVNCLDKFWHKACFHCETCKMTLNMKNYKGYEKKPYCNAHYPKQSFTMVADTPENLRLKQQSELQSQVRYKEEFEKNKGKGFSVVADTPELQRIKKTQDQISNIKYHEEFEKSRMGPSGGEGMEPERRDSQDGSSYRRPLEQQQPHHIPTSAPVYQQPQQQPVAQSYGGYKEPAAPVSIQRSAPGGGGKRYRAVYDYSAADEDEVSFQDGDTIVNVQQIDDGWMYGTVERTGDTGMLPANYVEAI

**>Hs lasp-2** 270aa

MNPQCARCGKVVYPTEKVNCLDKYWHKGCFHCEVCKMALNMNNYKGYEKKPYCNAHYPKQSFTTVADTPENLRLKQQSELQSQVKYKRDFEESKGRGFSIVTDTPELQRLKRTQEQISN**VKYHEDFEKTKGRGFTPVVDDPVTERVRKNTQVVSDAAYKGVHPHIVEMDRRPGIIVAPVLPGAYQQSHSQGYGYMHQTSVSSMRSMQHSPNLRTYRAMYDYSAQDEDEVSFRDGDYIVNVQPIDDGWMYGTVQRTGRTGMLPANYIEFVN**

**>Hs nebulette** 1014aa

MRVPVFEDIKDETEEEKIGEEENEEDQVFYKPVIEDLSMELARKCTELISDIRYKEEFKKSKDKCTFVTDSPMLNHVKNIGAFISEAKYKGTIKADLSNSLYKRMPATIDSVFAGEVTQLQSEVAYKQKHDAAKGFSDYAHMKEPPEVKHAMEVNKHQSNISYRKDVQDTHTYSAELDRPDIKMATQISKIISNAEYKKGQGIMNKEPAVIGRPDFEHAVEASKLSSQIKYKEKFDNEMKDKKHHYNPLESASFRQNQLAATLASNVKYKKDIQNMHDPVSDLPNLLFLDHVLKASKMLSGREYKKLFEENKGMYHFDADAVEHLHHKGNAVLQSQVKYKEEYEKNKGKPMLEFVETPSYQASKEAQKMQSEKVYKEDFEKEIKGRSSLDLDKTPEFLHVKYITNLLREKEYKKDLENEIKGKGMELNSEVLDIQRAKRASEMASEKEYKKDLESIIKGKGMQAGTDTLEMQHAKKAAEIASEKDYKRDLETEIKGKGMQVSTDTLDVQRAKKASEMASQKQYKKDLENEIKGKGMQVSMDIPDILRAKRTSEIYSQRKYKDEAEKMLSNYSTIADTPEIQRIKTTQQNISAVFYKKEVGAGTAVKDSPEIERVKKNQQNISSVKYKEEIKHATAISDPPELKRVKENQKNISNLQYKEQNYKATPVSMTPEIERVRRNQEQLSAVKYKGELQRGTAISDPPELKRAKENQKNISNVYYRGQLGRATTLSVTPEMERVKKNQENISS**VKYTQDHKQMKGRPSLILDTPAMRHVKEAQNHISM**VKYHEDFEKTKGRGFTPVVDDPVTERVRKNTQVVSDAAYKGVHPHIVEMDRRPGIIVDLKVWRTDPGSIFDLDPLEDNIQSRSLHMLSEKASHYRRHWSRSHSSSTFGTGLGDDRSEISEIYPSFSCCSEVTRPSDEG**APVLPGAYQQSHSQGYGYMHQTSVSSMRSMQHSPNLRTYRAMYDYSAQDEDEVSFRDGDYIVNVQPIDDGWMYGTVQRTGRTGMLPANYIEFVN**

**>Xt lasp-1** 258aa

MNPSCARCGKIVYPTEKVNCLDKFWHKACFHCEICKMTLNMKNYKGYDKKPYCNAHYPKQSFTTVADTPENLRLKQQSELQSQVRYKEDFEKNKGKGFSVVADTPELQRIKKTQDQISNIKYHEEFDKSRMGNPLVDGDDFDRRGSDDGANYRRSSQSTPQQHHPPPASSSAYPQHQPQPKYGYQEPAAPVSSQRSAPAGGKRYRAVYDYNAADEDEVSFQDGDTIINVQQIDDGWMYGTVERTGDTGMLPANYVEAM

**>Xt lasp-2** 276aa

MNPQCARCGRVVYPTEKVNCLDKFWHKGCFQCEVCKMTLNMNNYKGYDKKPYCNAHYPKQSFTTVTDTPENLRLKQQSEIQSQVKYKKDFEESKGRGFSIVTDALELQRLKKTQEQISN**IKYHEDFEKTKGKGFTPVVDDPITDRVRRNTQIVSDAAYRNVHPHVVEMDRRPGMVVGQGMPGAYPQHHPGYGYMHPSSAASSVRSAQPQHAPPLSPSMRTYRAMYDYCAQDEDEVSFREGDYIVNVQPIDEGWMYGTVQRSGKNGMLPANYIELVN**

**>Xt nebulette** 934aa

MKVPVIIDSTVDDSDEEEKVFPKPVIEDRNIEFAKKCTELISDVTYKEQYEKSKGKCAFVPDTLTLKNVSDSISETKYKEAGKKGLSNSLYKMMPATIDTAFAKEVSKLQSEVLYRQKYDAEKGKSNYAQMQEPPDVKHAKEVNKHQSNVSYKKDVLDIHKYTEVSDRLDIQKATEAAKFISNIEYKKGRENMNKEPVVLGRPDFEHAKEVSKLSSQVKYKEKFDKEMKGKKPCYNPLDSASFKQAQAANALASNVKYRQHLQDIHDPSVDMPNLLYLQHALKTSKMQSNNMYKKKFEEQKGIYHFALDTAEQIHHKENAVLQSQVKYKEEYEKSKGRSLLEFVDTPSYQVSKEAQKIQSEKEYRKDLEEGIKGKGLTVLEDTPDLIRVKNAAQILNEKEYRKDLENDIKGKGMEFSADIPDMIRAKRASEIASQKEYTKDYETEIKGKGMIIGTDIPEIQRVKRVTEMVSQKGYKKDLESEIKGKGMQIGADMPEIQRVKRVSEIVSQKEYTKDLEREIRGKGMQVGTDIPEIQRVKRVSEMVSQKEYKKDLETEIKGKGMQVGPDIPDLQRAMKASEIASQKKYKDEAEKLLSTYSIIADTPEIARVKATQKNISSVLYKKGVGAGTAVKDTPETERVKISQQNISAVKYKDDIGPATTISDLPEVKRVKEIQKNISNVQYKEQQLKATPVSVTPEMERVKKNQEHFSTVHYKKKVGKATAVSYTPEMERVKKNQENISSVKYSNDLKQLKGRPSVILDTPEMRHVKETQNHISM**IKYHEDFEKTKGKGFTPVVDDPITDRVRRNTQIVSDAAYRNVHPHVVEMDRRPGMVVGQGMPGAYPQHHPGYGYMHPSSAASSVRSAQPQHAPPLSPSMRTYRAMYDYCAQDEDEVSFREGDYIVNVQPIDEGWMYGTVQRSGKNGMLPANYIELVN**
